## Additional File 2 for "Choice of pre-processing pipeline influences clustering quality of scRNA-seq datasets"

### Zebrafish pineal gland scSeq sample 1 downstream analysis

Analysis of zebrafish pineal gland cell types. For original data see Shainer et al. 2019 (https://www.sciencedirect.com/science/article/pii/S0960982219305561 (https://www.sciencedirect.com/science/article/pii/S0960982219305561), pineal sample 1).

```
library(dplyr)
library(Seurat)
library(Matrix)
library(ggplot2)
library(reticulate)
library(stringr)
library(ggpubr)

#load files sample 1
pineal_s1_cellr_101 <- readRDS("/zstorage/cellr_vs_kallisto/rds_files/D_rerio.GRCz11.101/dr_pine
al_s1_cellr.rds")
pineal_s1_kb_101 <- readRDS("/zstorage/cellr_vs_kallisto/rds_files/D_rerio.GRCz11.101/dr_pineal_
s1_kb.rds")
pineal_s1_kb_forced_101 <- readRDS("/zstorage/cellr_vs_kallisto/rds_files/D_rerio.GRCz11.101/dr_
pineal_s1_kb_forced.rds")
```

### Downstream analysis of data preprocessed with kallisto

Calculate the percentage of mitochondrial genes per cell.

```
pineal_s1_kb_101[["percent.mt"]] <- PercentageFeatureSet(object = pineal_s1_kb_101, pattern = "^
mt-")
```

Visualize QC metrics.

```
VlnPlot(object = pineal_s1_kb_101,
        features = c("nFeature_RNA", "nCount_RNA", "percent.mt"),
        ncol = 3,
        pt.size=0)
```

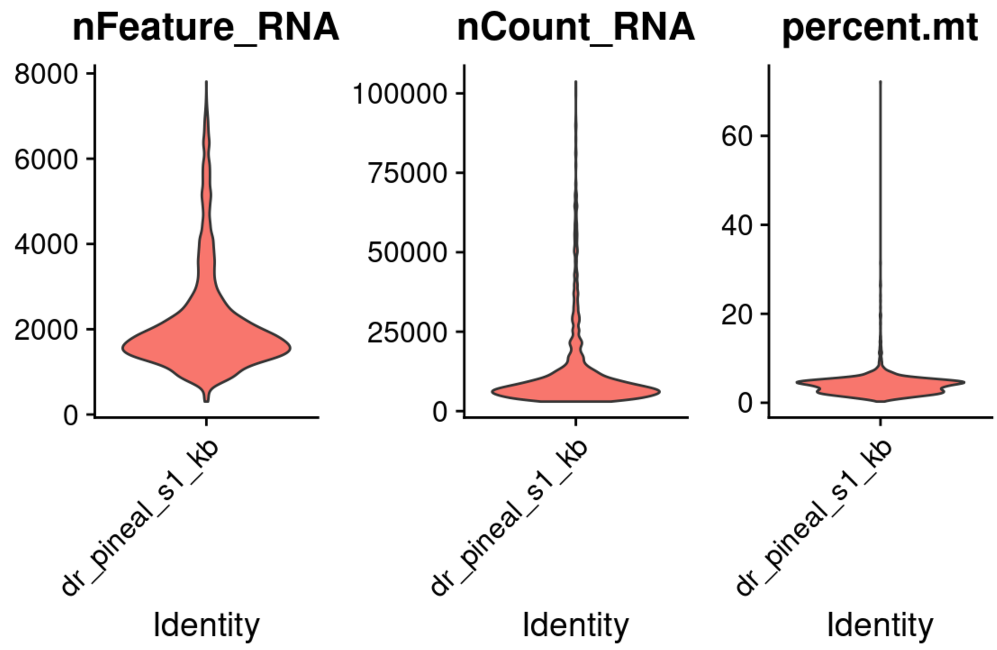


Total number of cells before filtration:

```
sum(table(...=pineal_s1_kb_101@active.ident))
```

```
## [1] 2675
```

Filtration of outlier cells containing unusual number of genes, UMI or percentage of mitochondrial genes. Plot the distribution of the filtered cells.

```
pineal_s1_kb_101 <- subset(x = pineal_s1_kb_101,
                           subset = nFeature_RNA > 200
                           & nCount_RNA < 15000
                           & percent.mt<30)

VlnPlot(object = pineal_s1_kb_101,
        features = c("nFeature_RNA", "nCount_RNA", "percent.mt"),
        ncol = 3,
        pt.size=0)
```

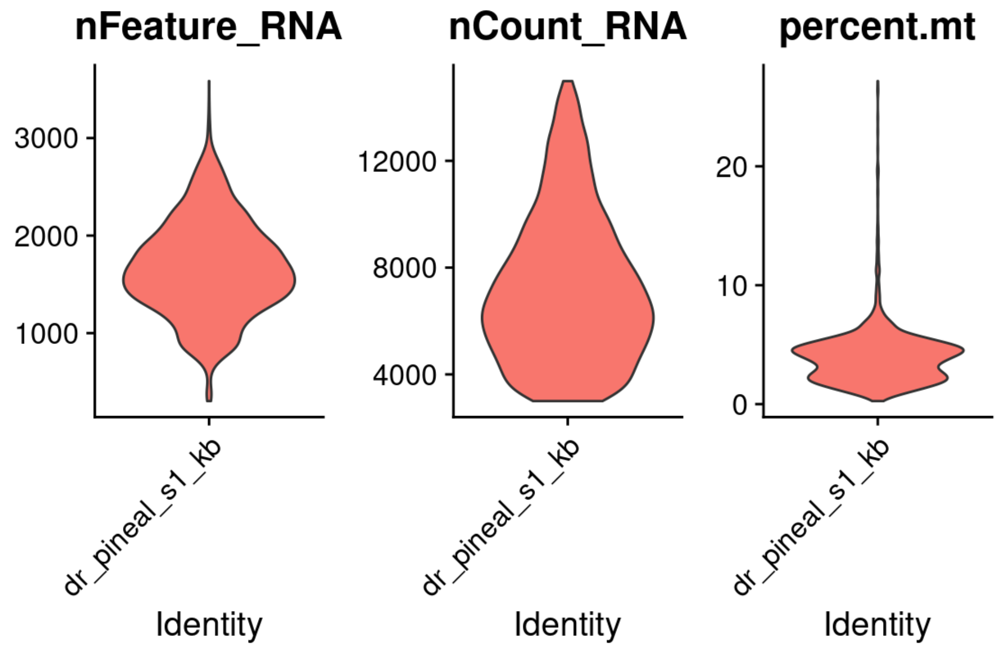

Total number of cells after filtration:

```
sum(table(...=pineal_s1_kb_101@active.ident))
```

```
## [1] 2199
```

Standard normalization, variable gene identification and scaling:

```
pineal_s1_kb_101 <- NormalizeData(object = pineal_s1_kb_101,
                                  normalization.method = "LogNormalize",
                                  scale.factor = 10000)

pineal_s1_kb_101 <- FindVariableFeatures(object = pineal_s1_kb_101,
                                         selection.method = "vst",
                                         nfeatures = 2000)

all_genes_kallisto_s1 <- rownames(x = pineal_s1_kb_101)
pineal_s1_kb_101 <- ScaleData(object = pineal_s1_kb_101, features = all_genes_kallisto_s1)
```

principal component analysis.

```
pineal_s1_kb_101 <- RunPCA(object = pineal_s1_kb_101, features = VariableFeatures(object = pinea
l_s1_kb_101))
```

```
## PC_ 1
## Positive:  selenop, krt8, actb2, atp1b1a, cd63, zgc:162730, actb1, s100a10b, junba, cxcl18b
##     atp1a1b, ctsla, icn, si:ch73-335l21.4, CR318588.4, zfp36l1b, sparc, fabp7a, krt18a.1, npc
2
##     fosl1a, tagln2, btg2, sgk1, fxyd1, nfkbiaa, gstp1, glulb, fosb, angptl4
## Negative:  rbp4l, tph2, pde6ha, rcvrn2, gngt1, pde6ga, exorh, saga, rcvrn3, arl3l1
##     gnat1, rbp3, BX004774.2, nrgna, tulp1b, cetn4, arl3l2, tulp1a, pde6gb, ahcy
##     asmt, si:dkey-220f10.4, cnga1a, ddc, unc119.2, gch1, cldn2, PLEKHB1, cngb3.2, ppdpfa
## PC_ 2
## Positive:  clu, fabp7a, atp1a1b, hsp70l, vim, krt18a.1, stra6, bckdhbl, bag3, cdkn1a
##     ubb, hspb8, sparc, rpe65a, slc38a3a, btg2, scg3, s100b, wfdc2, proca1
##     cxcl18b, si:ch211-80h18.1, fabp7b, zgc:114181, rgrb, dkk3b, rbp1.1, slc13a1, si:ch211-81a
5.8, cbx7a
## Negative:  si:dkey-27i16.2, cx32.2, fcer1gl, ctss2.2, zgc:64051, ccl35.1, lgals3bpb, si:cabz0
1074946.1, cd74b, cd74a
##     mhc2a, spi1b, stoml3b, havcr1, fermt3b, ccl34b.1, slc7a7, ccl35.2, ncf1, apoc1
##     cxcr4b, BX649485.1, lgals9l1, ms4a17a.9, ctsc, cebpa, laptm5, grna, mpeg1.1, pfn1
## PC_ 3
## Positive:  si:ch211-81a5.8, zgc:114181, ndrg1a, bckdhbl, slc13a1, dkk3b, fabp7b, gstm.3, si:d
key-11c5.11, si:ch211-80h18.1
##     rbp1.1, si:dkey-12l12.1, vim, AL954359.2, tmem98, mstnb, zgc:153311, flr, zgc:158404, igf
bp2a
##     foxj1a, tmem72, tbata, bmp1b, mt2, mcl1a, ppdpfa, proca1, ackr3a, rbp2b
## Negative:  si:dkey-33c12.3, snap25a, chga, bdnf, elavl3, vgf, elavl4, anxa13l, id4, gng13b
##     stmn4l, pcp4a, si:dkeyp-69c1.7, ywhag2, rtn1b, uts2a, nell2b, nxph1, hpcal4, nrsn1
##     cart3, p2rx2, atp6v1b2, stmn2a, sncga, tmsb2, insm1b, atp1a3a, pam, scg2b
## PC_ 4
## Positive:  zgc:114181, dkk3b, slc13a1, fabp7b, igfbp2a, gstm.3, si:dkey-12l12.1, rbp1.1, mstn
b, ahcyl1
##     flr, rbp4, bckdhbl, si:dkey-11c5.11, b3glcta, AL954359.2, tmem98, zgc:153311, slc38a3a, s
i:ch211-80h18.1
##     ndrg1a, ackr3a, tmem72, si:ch211-81a5.8, C7H20orf27, bmp1b, tbata, zgc:158404, vim, enkur
## Negative:  rdh5, rgra, asip2b, rbp5, fxyd6l, si:ch211-251b21.1, rlbp1b, cxcl14, efhd1, syt5b
##     scg3, jhy, ppp1r14aa, slc3a2a, slc6a2, slc1a3a, cldn7a, her15.2, tagln2, fdx1
##     coch, prdx1, myl9b, lgals2a, bzw1b, c1qtnf5, rpe65a, cnn2, cdon, txn
## PC_ 5
## Positive:  dcn, CABZ01092746.1, ccl25b, pmp22a, cldn11a, sost, fsta, twist1a, pcolcea, msx1b
##     lxn, thbs4b, anxa1a, bhmt, si:ch1073-291c23.2, clec3ba, cygb1, tmem176, selenbp1, eif4ebp
3
##     srgn, igfbp5b, si:dkey-11f4.20, zgc:158343, fstb, si:ch73-86n18.1, crhbp, adh8a, cxcl12a,
serpinf1
## Negative:  ctsd, stra6, scg3, rpe65a, fabp7a, lgals2a, hspb8, rgrb, atp1b1a, nanos1
##     efhd1, rgra, rdh5, rlbp1b, clu, si:dkey-112a7.4, actb1, si:ch211-251b21.1, jhy, stmn1b
##     bag3, syt5b, rnaseka, keap1a, smox, fdx1, cdkn1a, itm2cb, asip2b, uchl1
```

Visualize the principal components percentage of variance by an elbow plot.

```
ElbowPlot(object = pineal_s1_kb_101, ndims = 30)
```

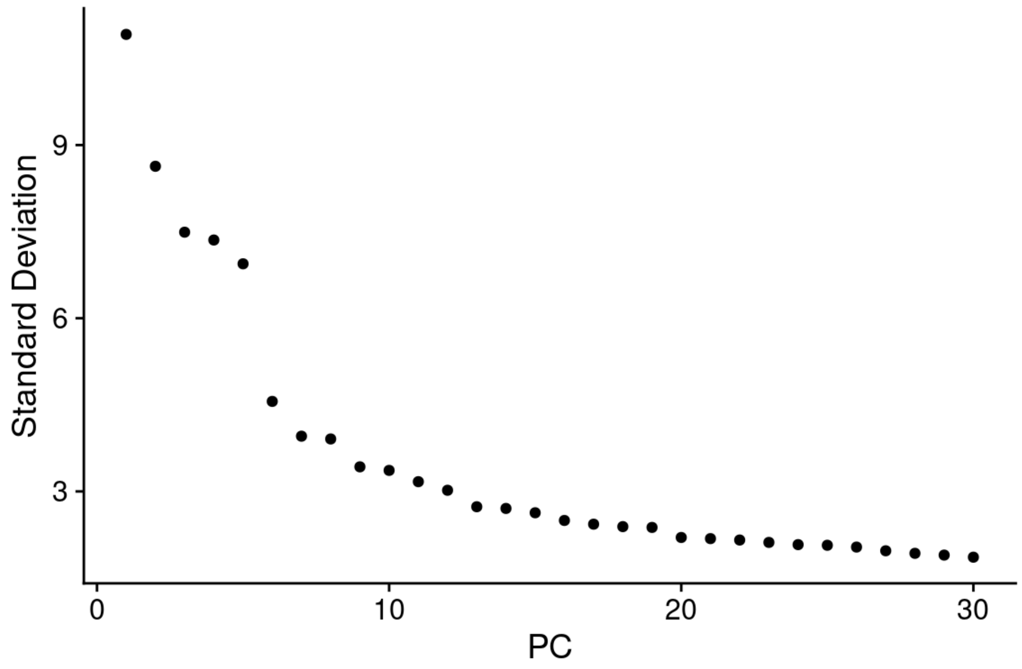


PCs 1-25 were used as dimensions of reduction to compute the k.param nearest neighbors

```
pineal_s1_kb_101 <- FindNeighbors(object = pineal_s1_kb_101, dims = 1:25)
pineal_s1_kb_101 <- FindClusters(object = pineal_s1_kb_101, resolution = 1.5)
```

```
## Modularity Optimizer version 1.3.0 by Ludo Waltman and Nees Jan van Eck
##
## Number of nodes: 2199
## Number of edges: 72904
##
## Running Louvain algorithm...
## Maximum modularity in 10 random starts: 0.8244
## Number of communities: 20
## Elapsed time: 0 seconds
```

```
pineal_s1_kb_101 <- RunUMAP(object = pineal_s1_kb_101, dims = 1:25)

kb_UMAP_unmerged_s1 <- DimPlot(object = pineal_s1_kb_101, reduction = "umap",
                            label=TRUE, pt.size = 0.5, label.size = 3) +
  theme(legend.position="none",
        axis.title.x=element_text(size=12),
        axis.title.y=element_text(size=12),
        plot.title = element_text(size=14, hjust=0.0)) + ggtitle("kallisto (res.=1.5)") +
  theme(plot.title = element_text(size = 12))
kb_UMAP_unmerged_s1
```

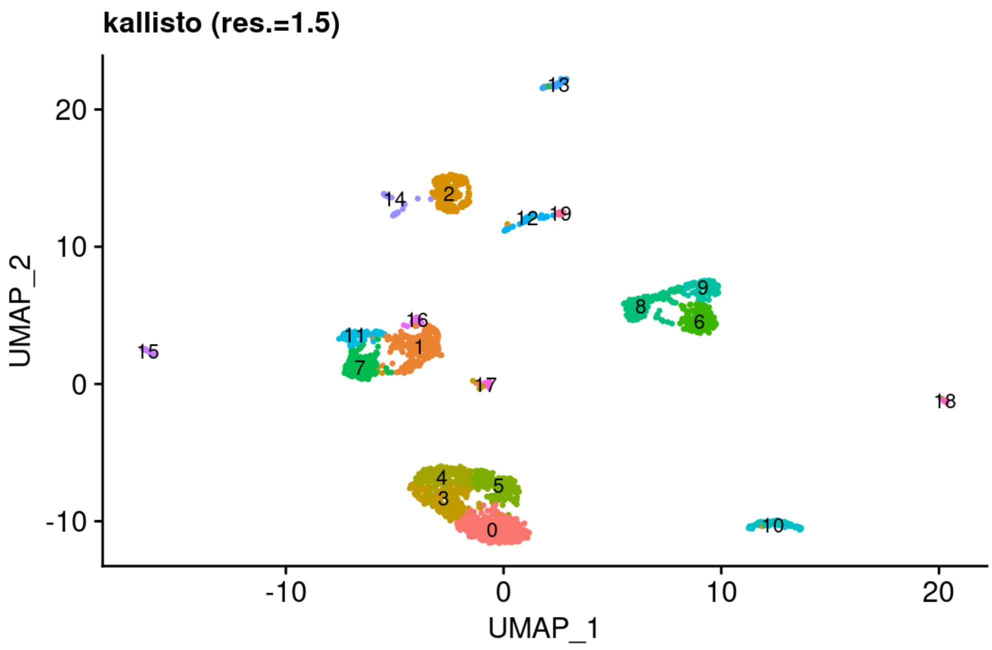


Analysis of the top markers for each cluster.

```
pineal_s1_kb_101.markers <- FindAllMarkers(object = pineal_s1_kb_101,
                                           only.pos = TRUE,
                                           min.pct = 0.25,
                                           logfc.threshold = 0.8)
```

```
pineal_s1_kb_101.markers %>% group_by(cluster) %>% top_n(n = 10, wt = avg_log2FC)
```

| p_val <dbl> | avg_log2FC <dbl> | pct.1 <dbl> | pct.2 <dbl> | p_val_adj <dbl> | cluster <fct> | gene <chr> |
|---|---|---|---|---|---|---|
| 3.165005e-163 | 2.071715 | 0.985 | 0.320 | 7.008270e-159 | 0 | cnga1a |
| 1.692290e-161 | 2.389093 | 1.000 | 0.903 | 3.747238e-157 | 0 | gngt1 |
| 4.744876e-155 | 1.859762 | 0.959 | 0.288 | 1.050658e-150 | 0 | CABZ01073265.1 |
| 6.156862e-153 | 2.206997 | 1.000 | 0.532 | 1.363314e-148 | 0 | saga |
| 9.347114e-150 | 2.005046 | 1.000 | 0.592 | 2.069731e-145 | 0 | unc119.2 |
| 5.523012e-142 | 2.324616 | 1.000 | 0.753 | 1.222961e-137 | 0 | exorh |
| 8.368707e-142 | 1.925816 | 1.000 | 0.438 | 1.853083e-137 | 0 | rbp3 |
| 4.763039e-137 | 1.994278 | 1.000 | 0.504 | 1.054680e-132 | 0 | gnat1 |
| 5.992704e-136 | 1.920668 | 1.000 | 0.517 | 1.326964e-131 | 0 | nrgna |
| 1.024534e-134 | 2.096840 | 1.000 | 0.990 | 2.268626e-130 | 0 | rbp4l |

1-10 of 200 rows Previous **1** 2 3 4 5 6 ... 20 Next

Dotplot of the top known markers of the pineal cell types (based on Shainer et al. 2019) as well as newly identify markers (such as col14a1b, dcn and ccr9a).

```
dot_plot_genes_s1= c("exorh", "gnat1", "gngt1",
                  "gnat2", "gngt2a", "col14a1b", "opn1lw1","parietopsin",
                  "asip2b", "rpe65a",
                  "dkk3b", "fabp7b",
                  "elavl4", "cart3",
                  "dcn", "igfbp5b",
                  "cahz", "hbaa1",
                  "ccr9a", "il4",
                  "cd74a", "apoc1",
                  "kdrl", "plvapa",
                  "bscl2l", "gch2",
                  "hbegfb", "fgl2a")

kallisto_dotplot_unmerged_s1<- DotPlot(pineal_s1_kb_101, features = dot_plot_genes_s1,
                                          cluster.idents=FALSE, dot.scale=2) + RotatedAxis() +
  theme(axis.text.x = element_text(angle=45, size=10),
        axis.text.y = element_text(size=5, angle=0),
        legend.title = element_text(size=10),
        legend.text = element_text(size = 10),
        axis.title.x=element_blank(),
        axis.title.y=element_blank())
kallisto_dotplot_unmerged_s1
```

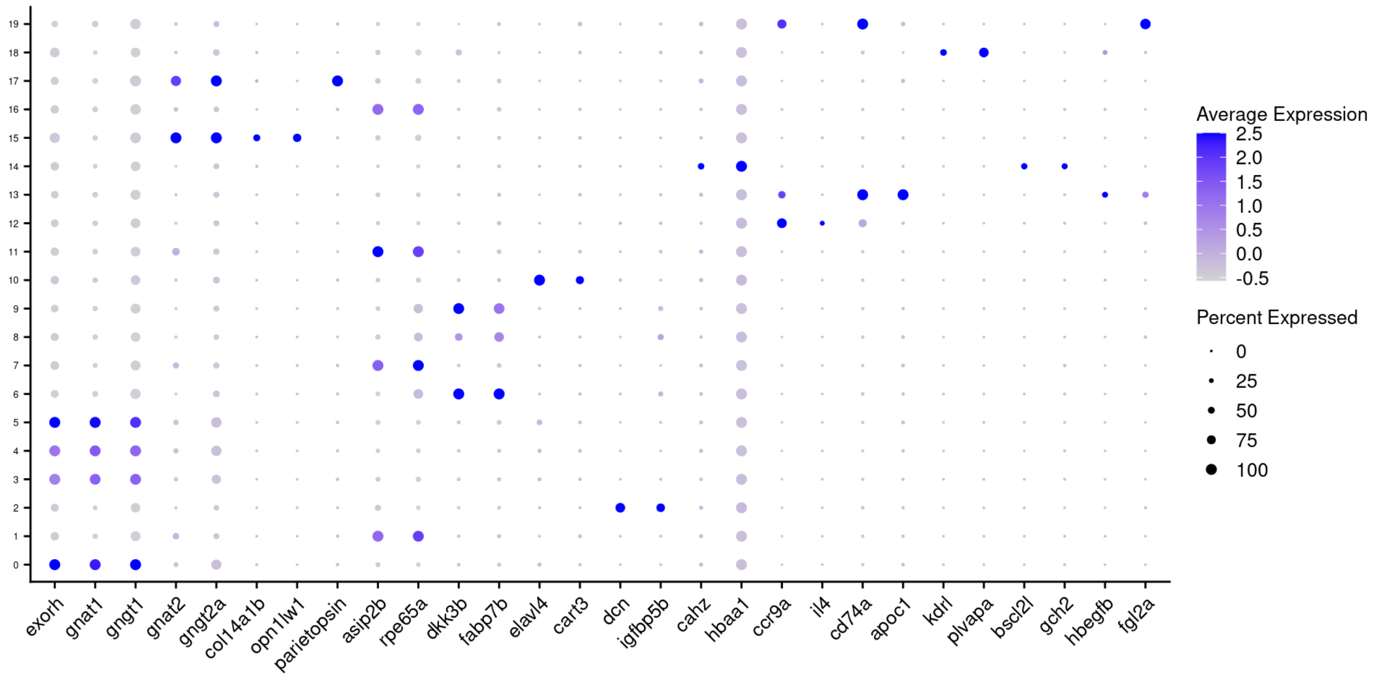


Markers separating the red- and green-like photoreceptors (clusters 15 & 17):

```
FeaturePlot(object=pineal_s1_kb_101, features = c("col14a1b", "parietopsin"), label = TRUE, labe
l.size = 3)
```

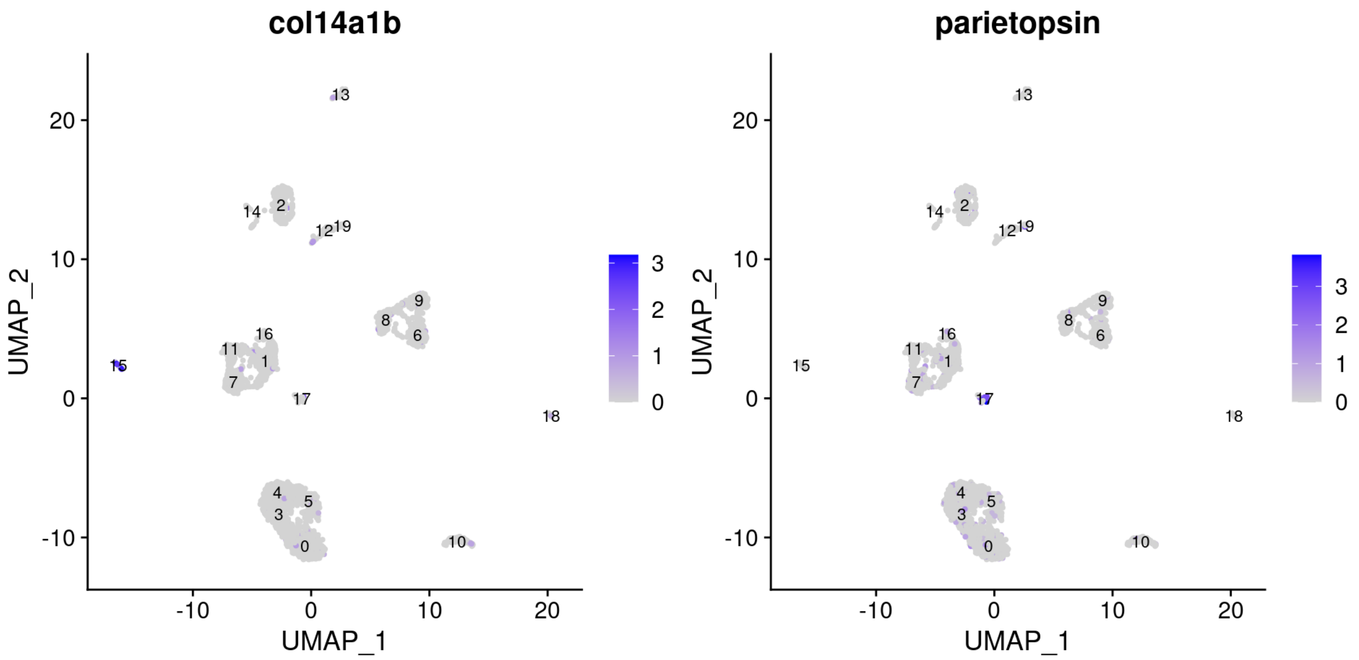


How many cells in each cluster?

```
(table(...=pineal_s1_kb_101@active.ident))
```

```
## ...
##   0   1   2   3   4   5   6   7   8   9  10  11  12  13  14  15  16  17  18  19
## 339 224 186 180 166 160 150 144 141  90  89  84  49  48  37  31  22  22  19  18
```

### Downstream analysis of data preprocessed with Cell Ranger

Calculate the percentage of mitochondrial genes per cell.

```
pineal_s1_cellr_101[["percent.mt"]] <- PercentageFeatureSet(object = pineal_s1_cellr_101, patter
n = "^mt-")
```

Visualize QC metrics.

```
VlnPlot(object = pineal_s1_cellr_101,
        features = c("nFeature_RNA", "nCount_RNA", "percent.mt"),
        ncol = 3,
        pt.size=0)
```

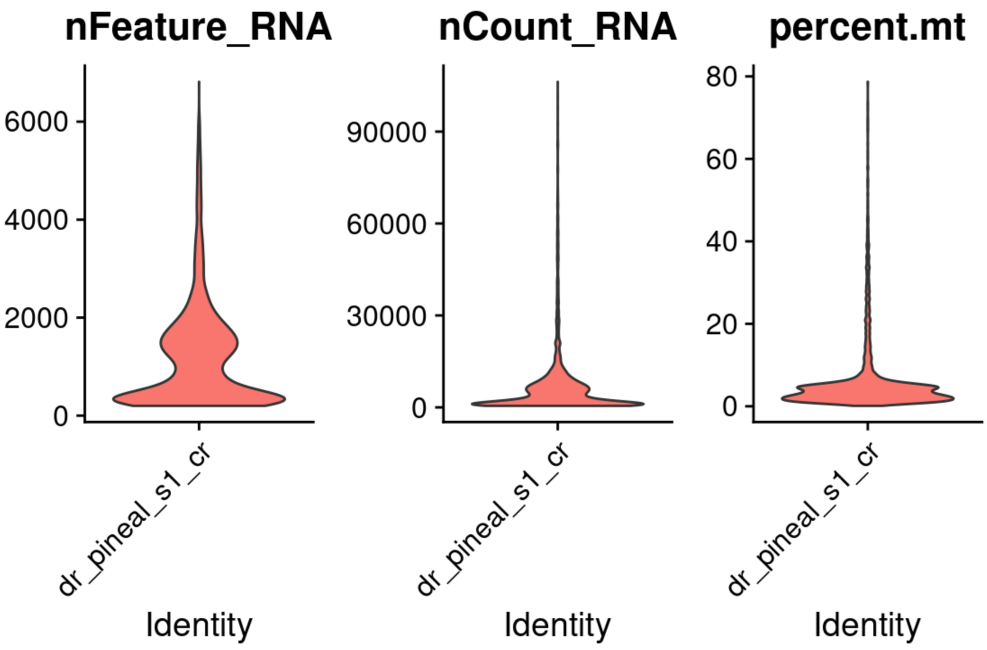


Total number of cells before filtration:

```
sum(table(...=pineal_s1_cellr_101@active.ident))
```

```
## [1] 4982
```

Filtration of outlier cells containing unusual number of genes, UMI or percentage of mitochondrial genes. Plot the distribution of the filtered cells.

```
pineal_s1_cellr_101 <- subset(x = pineal_s1_cellr_101,
                              subset = nFeature_RNA > 200
                              & nCount_RNA < 15000
                              & percent.mt<30)

VlnPlot(object = pineal_s1_cellr_101,
        features = c("nFeature_RNA", "nCount_RNA", "percent.mt"),
        ncol = 3,
        pt.size=0)
```

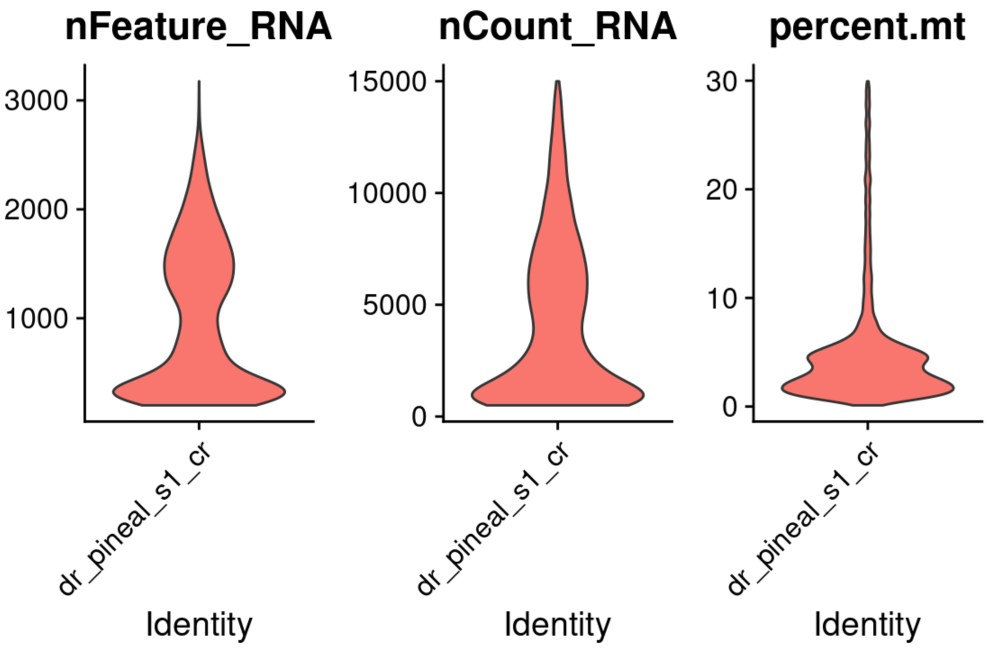

Total number of cells after filtration:

```
sum(table(...=pineal_s1_cellr_101@active.ident))
```

```
## [1] 4334
```

Standard normalization, variable gene identification and scaling:

```
pineal_s1_cellr_101 <- NormalizeData(object = pineal_s1_cellr_101,
                                  normalization.method = "LogNormalize",
                                  scale.factor = 10000)

pineal_s1_cellr_101 <- FindVariableFeatures(object = pineal_s1_cellr_101,
                                         selection.method = "vst",
                                         nfeatures = 2000)

all_genes_cellr_s1 <- rownames(x = pineal_s1_cellr_101)
pineal_s1_cellr_101 <- ScaleData(object = pineal_s1_cellr_101, features = all_genes_cellr_s1)
```

Principal component analysis.

```
pineal_s1_cellr_101 <- RunPCA(object = pineal_s1_cellr_101,
                              features = VariableFeatures(object = pineal_s1_cellr_101))
```

```
## PC_ 1
## Positive:  rbp4l, pde6ha, exorh, gngt1, pde6ga, gnat1, saga, rbp3, pde6gb, tph2
##     BX004774.2, arl3l1, nrgna, rcvrn2, tulp1a, cnga1a, tulp1b, rcvrn3, pde6a, ddc
##     arl3l2, si:dkey-220f10.4, rcvrnb, asmt, aipl2, ppdpfa, cetn4, ahcy, guk1b, parapinopsinb
## Negative:  selenop, krt8, zgc:162730, cd63, atp1b1a, s100a10b, si:ch73-335l21.4, junba, actb
2, cxcl18b
##     atp1a1b, icn, actb1, zfp36l1b, sparc, fxyd1, sgk1, angptl4, nfkbiaa, krt18a.1
##     CR318588.4, jdp2b, btg2, fabp7a, tagln2, ccn1, ctsla, igfbp2a, fosb, serpinh1b
## PC_ 2
## Positive:  si:dkey-27i16.2, fcer1gl, spi1b, ctss2.2, cd74a, cd74b, zgc:64051, lgals3bpb, ccl3
5.1, stoml3b
##     havcr1, si:cabz01074946.1, mhc2a, cx32.2, slc7a7, cxcr4b, ncf1, pfn1, ms4a17a.9, ncf2
##     arpc1b, BX649485.1, ccl34b.1, ccl35.2, ctsc, laptm5, mpeg1.1, lgals9l1, apoc1, fermt3b
## Negative:  clu, atp1a1b, fabp7a, bckdhbl, vim, stra6, bag3, hspb8, krt18a.1, hsp70l
##     si:ch211-81a5.8, cdkn1a, slc38a3a, hspa8b, cbx7a, scg3, wfdc2, si:ch211-80h18.1, proca1,
ubb
##     zgc:114181, slc13a1, dkk3b, mstnb, AL954359.2, s100b, fabp7b, lpl, cxcl18b, btg2
## PC_ 3
## Positive:  si:dkey-33c12.3, chga, bdnf, elavl4, snap25a, vgf, elavl3, p2rx2, vav3b, anxa13l
##     id4, stmn2a, nxph1, nell2b, tcf7l2, gng13b, hpcal4, syt1a, atp6v1b2, rtn1b
##     ywhag2, nrsn1, scg2b, stmn4l, si:dkeyp-69c1.7, insm1b, atp1a3a, cart3, pam, map1b
## Negative:  ppdpfa, si:ch211-81a5.8, pde6gb, rbp4l, pde6ha, BX004774.2, ahcy, gngt1, zgc:11418
1, rbp3
##     bckdhbl, tulp1b, ndrg1a, dkk3b, nrgna, unc119.2, gnat1, slc13a1, exorh, pde6ga
##     cnga1a, arl3l2, tulp1a, saga, pla1a, fabp7b, si:dkey-11c5.11, cetn4, pde6a, arl3l1
## PC_ 4
## Positive:  hbaa1, hbba1.1, hbba1, hbba2, si:ch211-5k11.8, ccl25b, hbaa2, dcn, CABZ01092746.1,
pmp22a
##     sost, crhbp, cldn11a, fsta, si:dkey-11f4.20, pcolcea, cxcl12a, lxn, cahz, thbs4b
##     twist1a, igfbp5b, bhmt, si:ch211-103n10.5, si:ch211-250g4.3, si:ch1073-291c23.2, msx1b, a
nxa1a, tmem176, eif4ebp3
## Negative:  ctsd, fkbp4, scg3, stoml3b, atp6v0ca, vmp1, uchl1, hspa8b, cct4, cx32.2
##     havcr1, lgals3bpb, CABZ01020840.1, spi1b, cbx7a, rdh5, slc7a7, ctss2.2, bag3, gnb1b
##     rnaseka, sqstm1, ctsc, hsp70l, atp1b1a, si:dkey-112a7.4, tspan36, stra6, sb:cb1058, fcer1
gl
## PC_ 5
## Positive:  asip2b, rdh5, rgra, fxyd6l, si:ch211-251b21.1, rbp5, efhd1, cxcl14, rlbp1b, syt5b
##     jhy, slc1a3a, her15.2, slc6a2, c1qtnf5, cldn7a, tagln2, slc3a2a, coch, ppp1r14aa
##     scg3, cdo1, marcksl1b, prdx1, fdx1, cnn2, bzw1b, myl9b, cdon, lgals2a
## Negative:  dkk3b, zgc:114181, slc13a1, fabp7b, pla1a, gstm.3, rbp1.1, bckdhbl, si:dkey-12l12.
1, AL954359.2
##     si:ch211-80h18.1, zgc:153311, mstnb, tmem98, si:dkey-11c5.11, cyp19a1b, slc38a3a, tmem72,
tbata, zgc:158404
##     slc13a3, b3glcta, vim, bmp1b, igfbp2a, rbp2b, ackr3a, C7H20orf27, ndrg1a, foxj1a
```

Visualize the principal components percentage of variance by an elbow plot.

```
ElbowPlot(object = pineal_s1_cellr_101, ndims = 30)
```

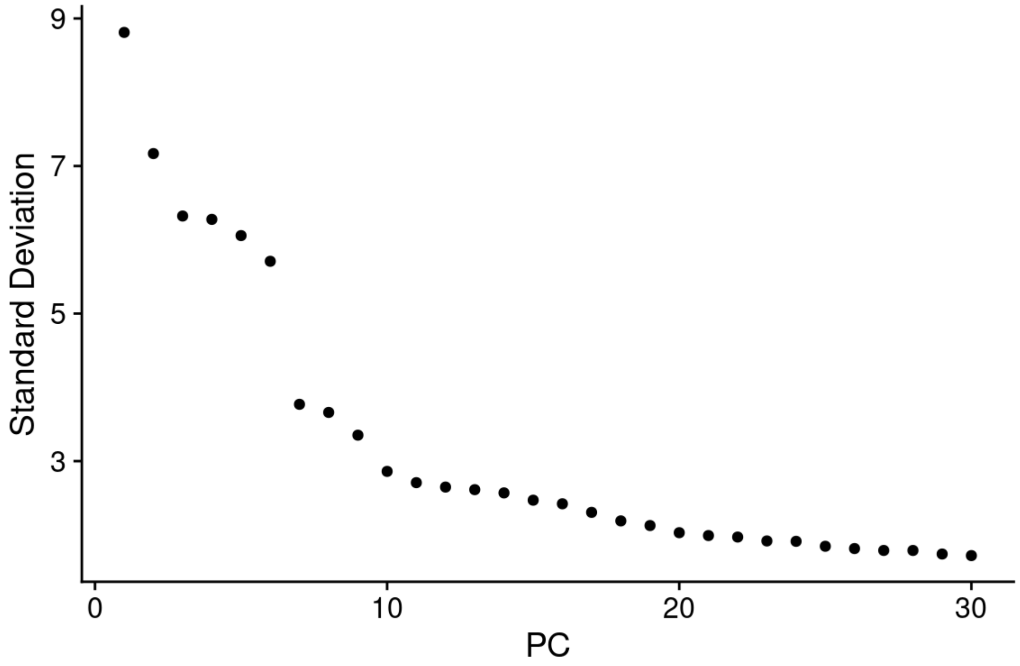


PCs 1-25 were used as dimensions of reduction to compute the k.param nearest neighbors

```
pineal_s1_cellr_101 <- FindNeighbors(object = pineal_s1_cellr_101, dims = 1:25)
pineal_s1_cellr_101 <- FindClusters(object = pineal_s1_cellr_101, resolution = 1.5)
```

```
## Modularity Optimizer version 1.3.0 by Ludo Waltman and Nees Jan van Eck
##
## Number of nodes: 4334
## Number of edges: 159944
##
## Running Louvain algorithm...
## Maximum modularity in 10 random starts: 0.8560
## Number of communities: 23
## Elapsed time: 0 seconds
```

```
pineal_s1_cellr_101 <- RunUMAP(object = pineal_s1_cellr_101, dims = 1:25)

cellr_UMAP_unmerged_s1 <- DimPlot(object = pineal_s1_cellr_101, reduction = "umap",
                            label=TRUE, pt.size = 0.5, label.size = 3) +
  theme(legend.position="none",
        axis.title.x=element_text(size=12),
        axis.title.y=element_text(size=12),
        plot.title = element_text(size=12, hjust=0.0)) + ggtitle("Cell Ranger (res.=1.5)")+
  theme(plot.title = element_text(size = 12))
cellr_UMAP_unmerged_s1
```

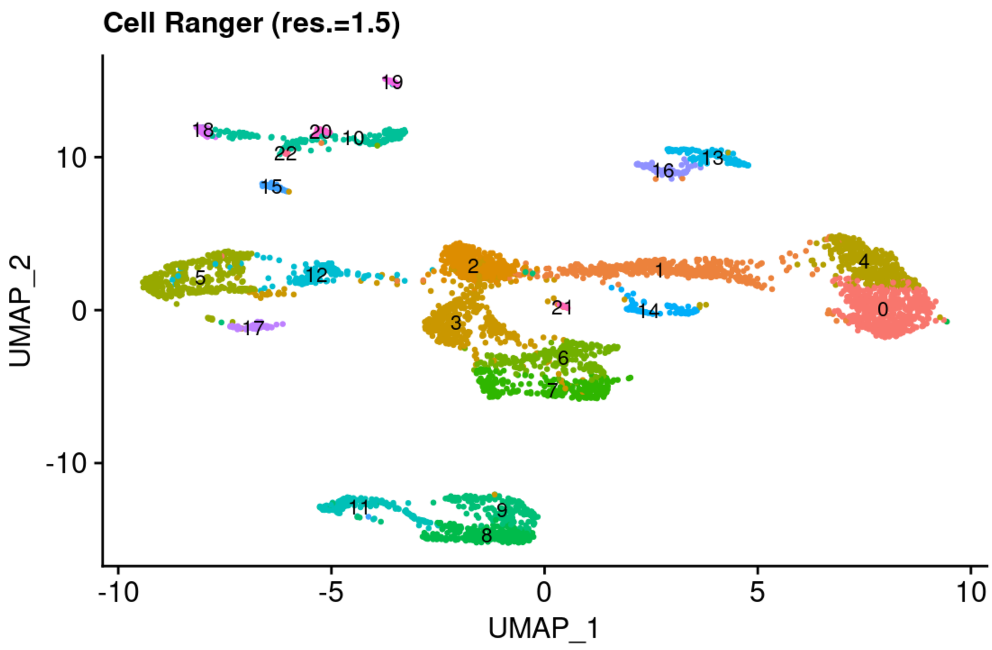


Analysis of the top markers for each cluster.

```
pineal_s1_cellr_101.markers <- FindAllMarkers(object = pineal_s1_cellr_101,
                                               only.pos = TRUE,
                                               min.pct = 0.25,
                                               logfc.threshold = 0.8)
```

```
pineal_s1_cellr_101.markers %>% group_by(cluster) %>% top_n(n = 10, wt = avg_log2FC)
```

| p_val <dbl> | avg_log2FC <dbl> | pct.1 <dbl> | pct.2 <dbl> | p_val_adj <dbl> | cluster <fct> | gene <chr> |
|---|---|---|---|---|---|---|
| 0.000000e+00 | 2.111182 | 0.994 | 0.205 | 0.000000e+00 | 0 | cnga1a |
| 1.158942e-282 | 2.577965 | 1.000 | 0.396 | 2.246493e-278 | 0 | gnat1 |
| 1.355631e-276 | 2.401663 | 1.000 | 0.382 | 2.627754e-272 | 0 | nrgna |
| 7.330473e-274 | 2.582430 | 1.000 | 0.435 | 1.420939e-269 | 0 | saga |
| 1.035648e-259 | 2.247651 | 1.000 | 0.413 | 2.007500e-255 | 0 | unc119.2 |
| 1.418734e-254 | 2.726601 | 1.000 | 0.855 | 2.750074e-250 | 0 | gngt1 |
| 9.601141e-250 | 2.174552 | 1.000 | 0.432 | 1.861085e-245 | 0 | ahcy |
| 9.072748e-248 | 2.667970 | 1.000 | 0.697 | 1.758662e-243 | 0 | exorh |
| 3.879519e-233 | 2.364702 | 1.000 | 0.876 | 7.520060e-229 | 0 | pde6gb |
| 2.882283e-214 | 2.207221 | 1.000 | 0.975 | 5.587018e-210 | 0 | rbp4l |

1-10 of 230 rows    Previous **1** 2 3 4 5 6 ... 23 Next

Dotplot of the top known markers of the pineal cell types (based on Shainer et al. 2019) as well as newly identify markers (such as dcn and ccr9a).

```
cellranger_dotplot_unmerged_s1<- DotPlot(pineal_s1_cellr_101, features = dot_plot_genes_s1,
                                          cluster.idents=FALSE, dot.scale=2) + RotatedAxis() +
  theme(axis.text.x = element_text(angle=45, size=10),
        axis.text.y = element_text(size=5, angle=0),
        legend.title = element_text(size=10),
        legend.text = element_text(size = 10),
        axis.title.x=element_blank(),
        axis.title.y=element_blank())
```

```
## Warning in FetchData(object = object, vars = features, cells = cells): The
## following requested variables were not found: col14a1b
```

```
cellranger_dotplot_unmerged_s1
```

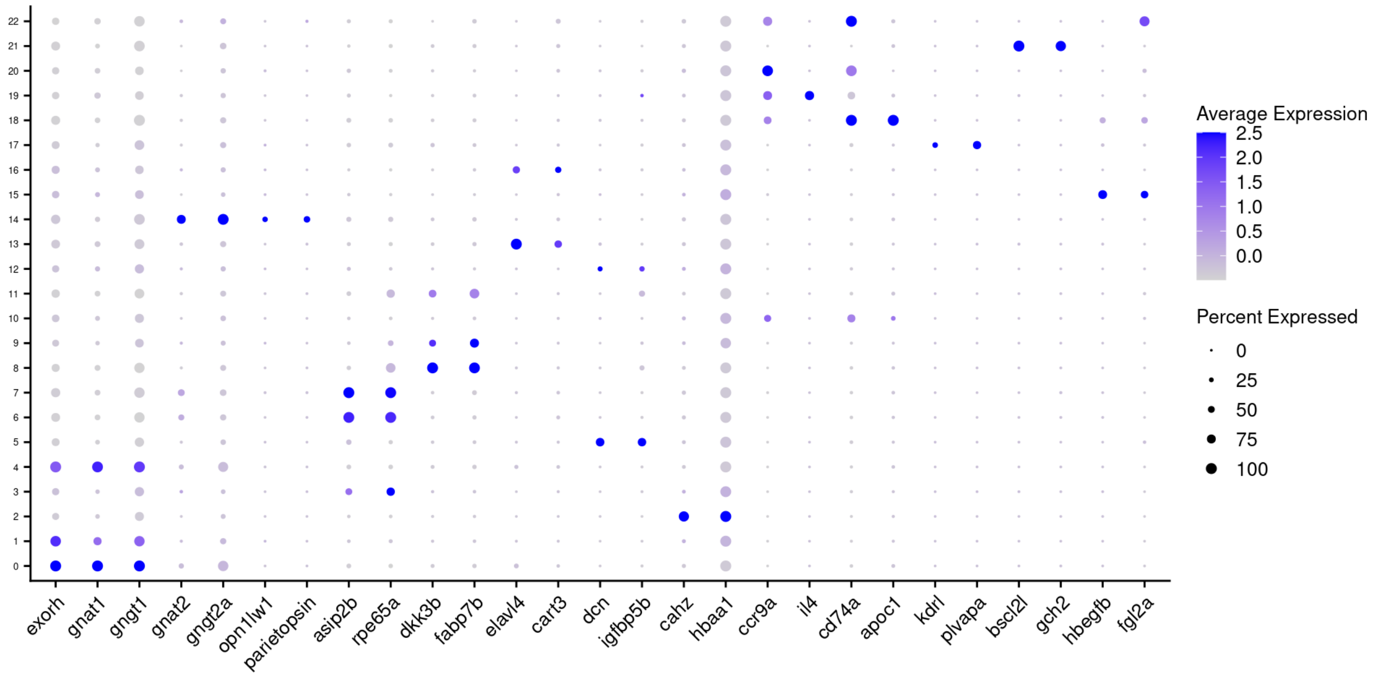


Markers for red- and green-like photoreceptors are expressed in a single cluster (#14):

```
FeaturePlot(object=pineal_s1_cellr_101, features = c("opn1lw1", "parietopsin"), label = TRUE, la
bel.size = 3)
```

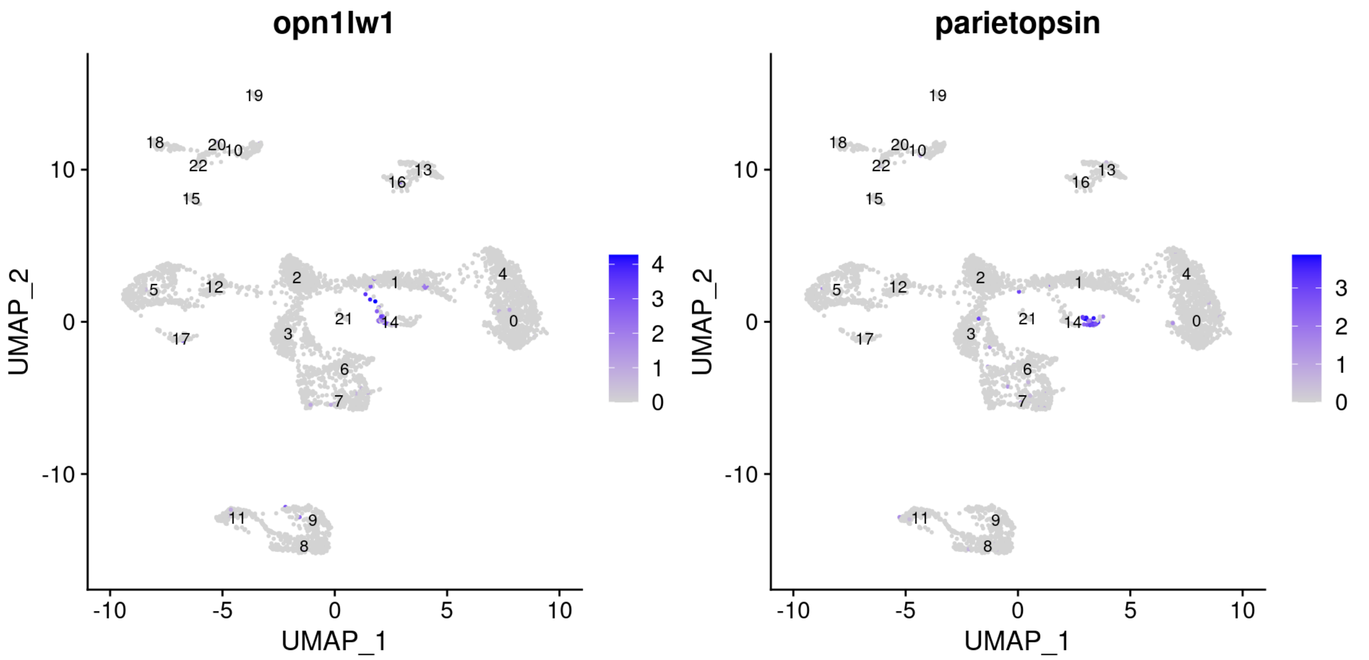


How many cells in each cluster?

```
(table(...=pineal_s1_cellr_101@active.ident))
```

```
## ...
##   0   1   2   3   4   5   6   7   8   9  10  11  12  13  14  15  16  17  18  19
## 485 421 412 403 364 364 262 253 240 174 159 154 127 112  80  62  60  51  46  32
##  20  21  22
##  31  24  18
```

When using the same parameters, the two types of cone-like photoreceptors can be distinguished when the pre-processing is done with kallisto, but not Cell Ranger, even though Kallisto pre-processed data contain half of the total cells (2199 cells in kallisto, and 4334 cells in Cell Ranger), and 2/3 of the cone-like photoreceptors (53 in kallisto and 80 in Cell Ranger).

Increasing the resolution does not improve the identification of the two cone-like photoreceptors types in the Cell Ranger data (the green and red opsin expressing cells are still considered the same cluster) :

```
pineal_s1_cellr_101 <- FindNeighbors(object = pineal_s1_cellr_101, dims = 1:25)
pineal_s1_cellr_101 <- FindClusters(object = pineal_s1_cellr_101, resolution = 3.5)
```

```
## Modularity Optimizer version 1.3.0 by Ludo Waltman and Nees Jan van Eck
##
## Number of nodes: 4334
## Number of edges: 159944
##
## Running Louvain algorithm...
## Maximum modularity in 10 random starts: 0.7604
## Number of communities: 38
## Elapsed time: 0 seconds
```

```
pineal_s1_cellr_101 <- RunUMAP(object = pineal_s1_cellr_101, dims = 1:25)

cellr_UMAP_unmerged_s1_res_3_5 <- DimPlot(object = pineal_s1_cellr_101, reduction = "umap",
                            label=TRUE, pt.size = 0.5, label.size = 3) +
  theme(legend.position="none",
        axis.title.x=element_text(size=10),
        axis.title.y=element_text(size=10),
        plot.title = element_text(size=12, hjust=0.0)) + ggtitle("Cell Ranger (res.=3.5)")
cellr_UMAP_unmerged_s1_res_3_5
```

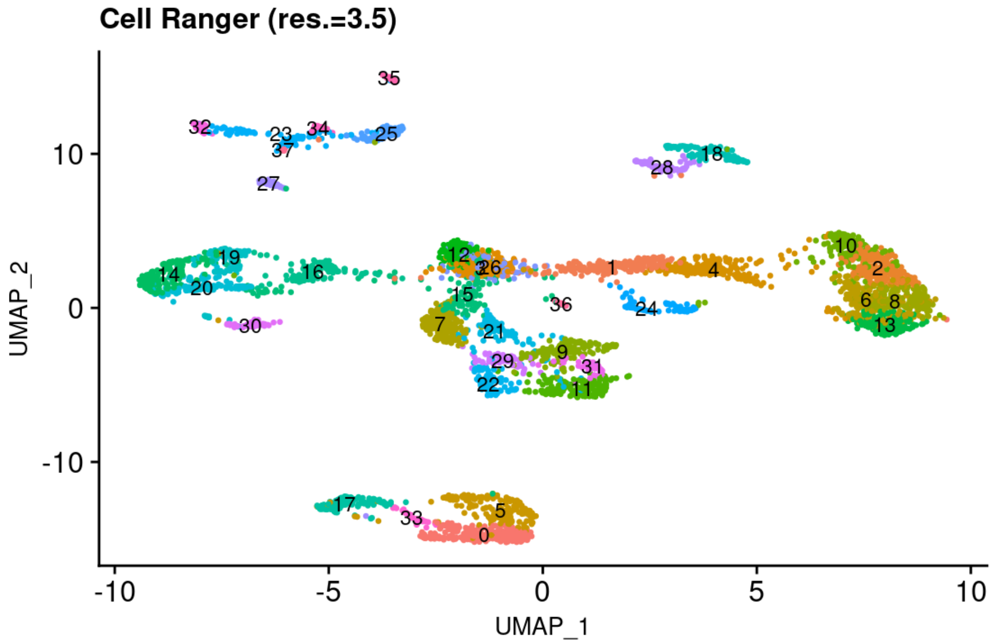


```
FeaturePlot(object=pineal_s1_cellr_101, features = c("opn1lw1", "parietopsin"), label = TRUE, la
bel.size = 3)
```

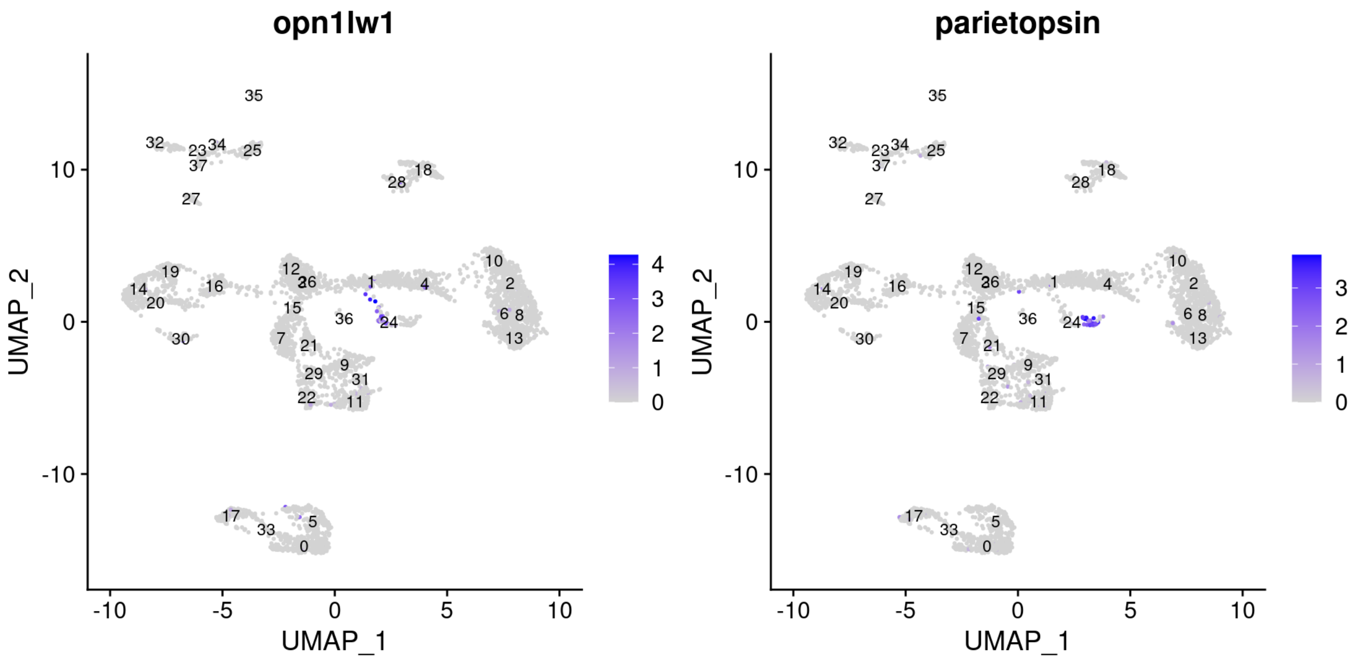


Dotplot high res.

```
cellranger_dotplot_high_res_s1<- DotPlot(pineal_s1_cellr_101, features = dot_plot_genes_s1,
                                         cluster.idents=FALSE, dot.scale=2) + RotatedAxis() +
  theme(axis.text.x = element_text(angle=45, size=10),
        axis.text.y = element_text(size=5, angle=0),
        legend.title = element_text(size=10),
        legend.text = element_text(size = 10),
        axis.title.x=element_blank(),
        axis.title.y=element_blank())
```

```
## Warning in FetchData(object = object, vars = features, cells = cells): The
## following requested variables were not found: col14a1b
```

```
cellranger_dotplot_high_res_s1
```

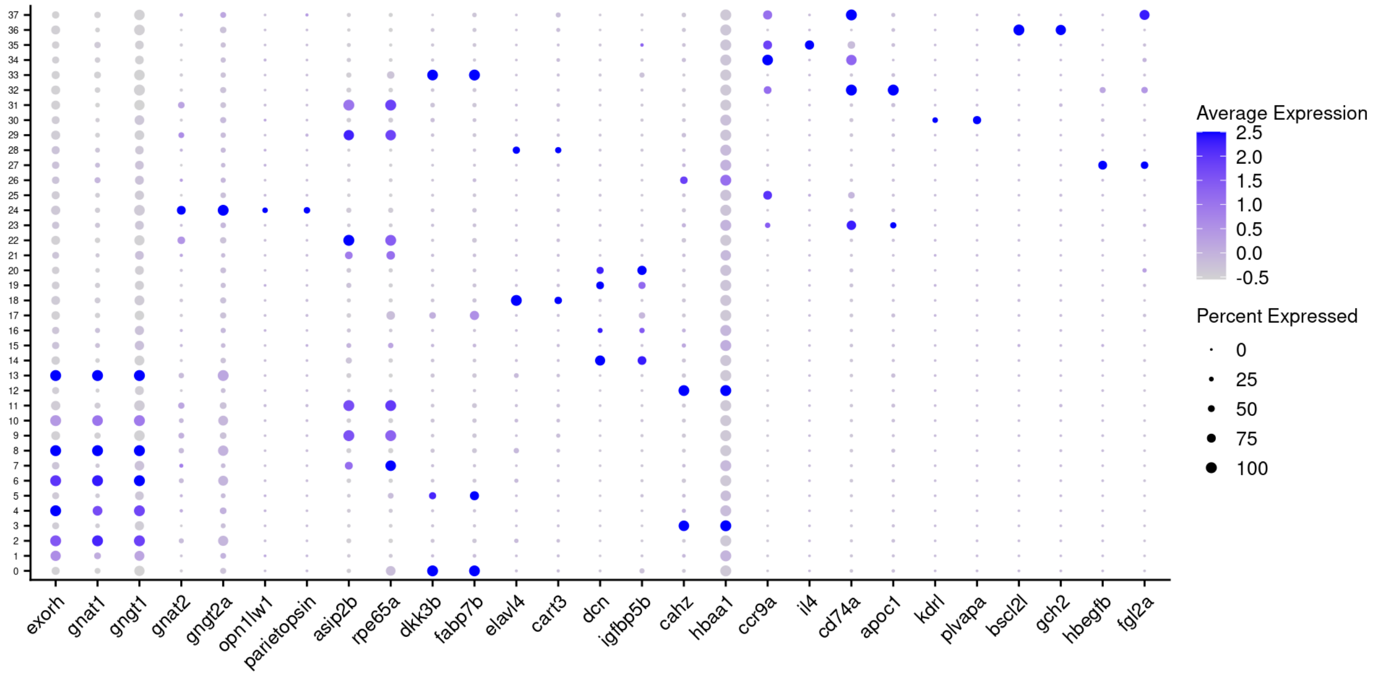


This shows that similar to the results observed for the pineal sample 2, the additional type of the photoreceptors can only be detected under standard conditions in the kallisto pre-processed data, but not for the Cell Ranger pre-processed data.

For sample number 1, additional differences were observed between the Cell Ranger and the kallisto pre-processed data. Two Cell Ranger clusters (#15 & #21, when analyzed with resolution of 1.5) did not express any of the described pineal markers (see dotplot). By exploring the markers of those cluster, we identified cluster 21 to be pigment cells (expressing bscl2l & gch2 genes, which are known xanthophore markers) and cluster 15 to be a type of hematopoietic cells (expressing hbegfb & fgl2a).

```
FeaturePlot(object=pineal_s1_cellr_101, features = c("bscl2l", "gch2", "hbegfb", "fgl2a"), label
= TRUE, label.size = 3)
```

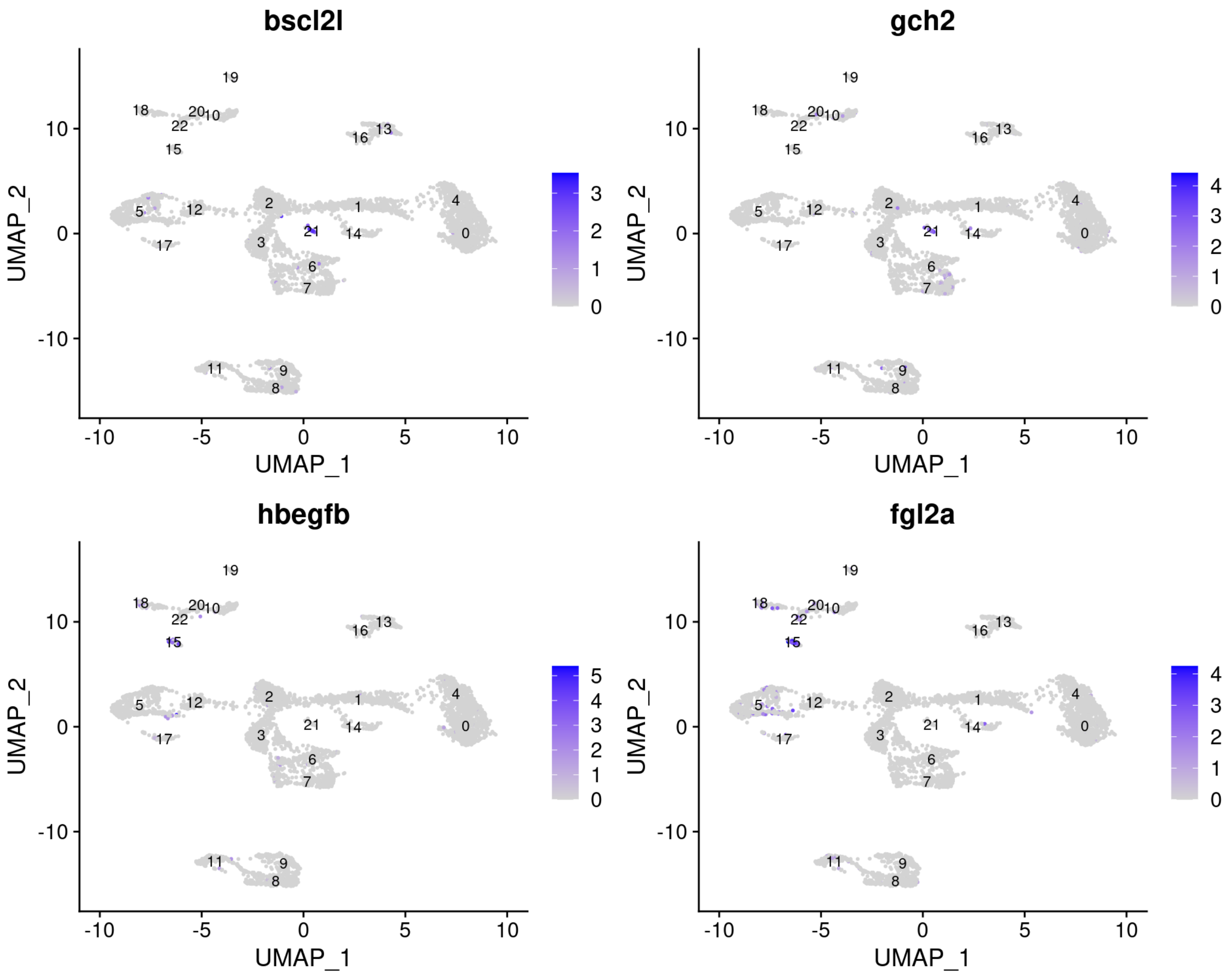


Although these are not true pineal specific cell type, but rather present a contamination of “outside cells” such as the pigment cells (originating from the skin), or hematopoietic cells that reside in any tissue, these clusters seem to be unique to the Cell Ranger dataset and do not appear in the kallisto data set. Are those cells appear in kallisto and assigned to another cluster or are they filtered out? To test this, we checked whether the cell barcodes of the cells belonging to cluster 15 and 21 in the Cell Ranger data exist in the kallisto data, and if so in which cluster:

```
# list all of kallisto cell barcodes
kb_cell_barcodes<-data.frame(WhichCells(pineal_s1_kb_101))
names(kb_cell_barcodes)[1]="barcodes"

# list Cell Ranger cluster #21 cell barcodes
cellr_21_barcodes<-data.frame(WhichCells(pineal_s1_cellr_101, idents = "21"))
names(cellr_21_barcodes)[1]="barcodes"
cellr_21_barcodes$barcodes<-str_remove(cellr_21_barcodes$barcodes, "[-1]")
cellr_21_barcodes$barcodes<-str_remove(cellr_21_barcodes$barcodes, "[1]")
#plot Cell Ranger cluster #21 cell barcodes that exist in kallisto (kallisto UMAP)
cluster_21_in_kb<-DimPlot(pineal_s1_kb_101,
        label=TRUE,
        cells.highlight = c(intersect(cellr_21_barcodes, kb_cell_barcodes)),
        label.size = 3) +
  ggtitle("Cell Ranger cluster #21 cell barcodes in kallisto") +
  theme(plot.title = element_text(size = 12, face="plain"))

# list Cell Ranger cluster #15 cell barcode
cellr_15_barcodes<-data.frame(WhichCells(pineal_s1_cellr_101, idents = "15"))
names(cellr_15_barcodes)[1]="barcodes"
cellr_15_barcodes$barcodes<-str_remove(cellr_15_barcodes$barcodes, "[-1]")
cellr_15_barcodes$barcodes<-str_remove(cellr_15_barcodes$barcodes, "[1]")
#plot Cell Ranger cluster #15 cell barcodes that exist in kallisto (kallisto UMAP)
cluster_15_in_kb<-DimPlot(pineal_s1_kb_101,
        label=TRUE,
        cells.highlight = c(intersect(cellr_15_barcodes, kb_cell_barcodes)),
        label.size = 3) +
  ggtitle("Cell Ranger cluster #15 cell barcodes in kallisto") +
  theme(plot.title = element_text(size = 12, face="plain"))

unique_barcodes_plot<-ggarrange(cluster_21_in_kb, cluster_15_in_kb,
          common.legend = TRUE,
          labels = c("A", "B"),
          ncol = 2, nrow = 1, legend = "right")
unique_barcodes_plot
```

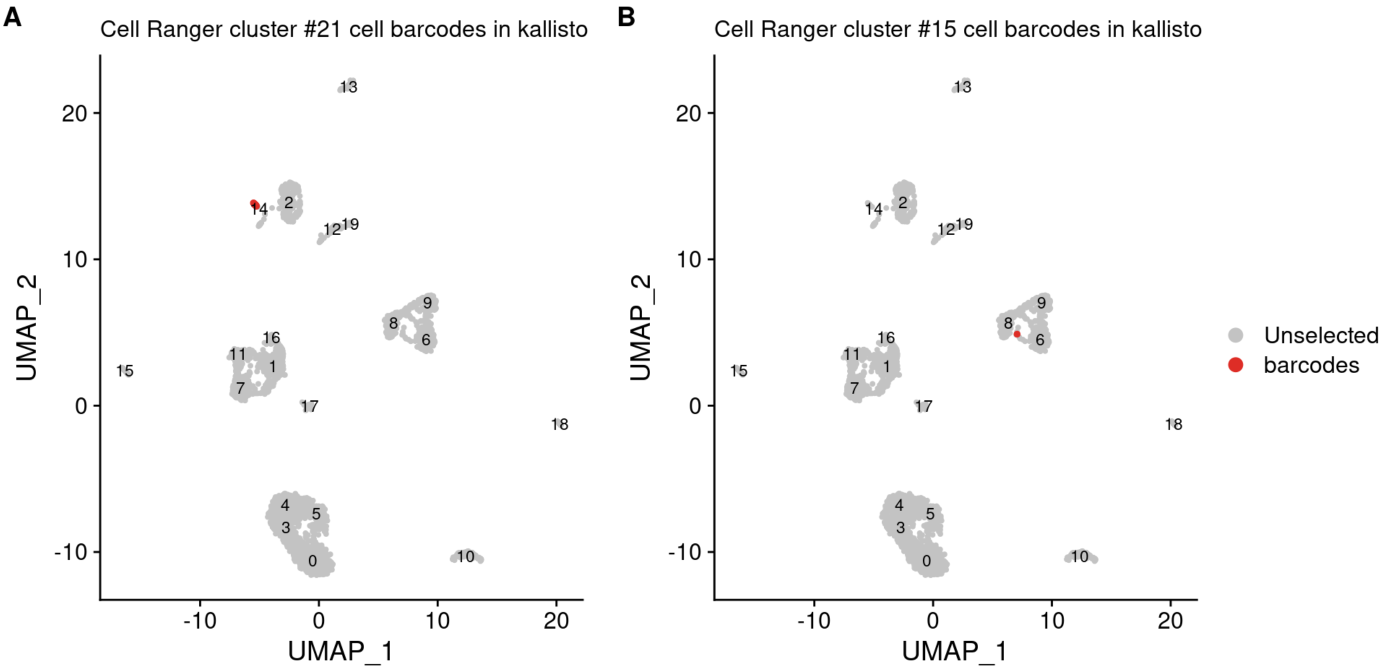


Cell Ranger cluster #15 has 62 cells and cluster #21 has 24 cells, the number of the same cells that appear in kallisto:

```
# The number of Cell Ranger cluster #15 cell barcodes that exist in kallisto:
nrow(intersect(cellr_15_barcodes, kb_cell_barcodes))
```

```
## [1] 1
```

```
# The number of Cell Ranger cluster #21 cell barcodes that exist in kallisto:
nrow(intersect(cellr_21_barcodes, kb_cell_barcodes))
```

```
## [1] 16
```

Cluster #15 cells are almost completely filtered out in the kallisto dataset. These cells might be of low quality or are just lost. Cluster #21 cells mostly exist in the kallisto (but in a lower number), but do not form a unique cluster under the conditions set for the clustering of the data (UMAP shows that this cluster can be assigned manually or by other clustering parameters, similar to the case of the green-like photoreceptors described in the manuscript), or as a result of the lower number of cells of this type. This shows that some clusters can be observed in the Cell Ranger pre-processed data that do not appear in the kallisto data. In this particular case, these clusters represent an "uninteresting" information, as those are not true pineal gland cell types, but for other datasets it might be of a biological relevance.

### Downstream analysis of data preprocessed with kallisto_forced

Calculate the percentage of mitochondrial genes per cell.

```
pineal_s1_kb_forced_101[["percent.mt"]] <- PercentageFeatureSet(object = pineal_s1_kb_forced_10
1, pattern = "^mt-")
```

Visualize QC metrics.

```
VlnPlot(object = pineal_s1_kb_forced_101,
        features = c("nFeature_RNA", "nCount_RNA", "percent.mt"),
        ncol = 3,
        pt.size=0)
```

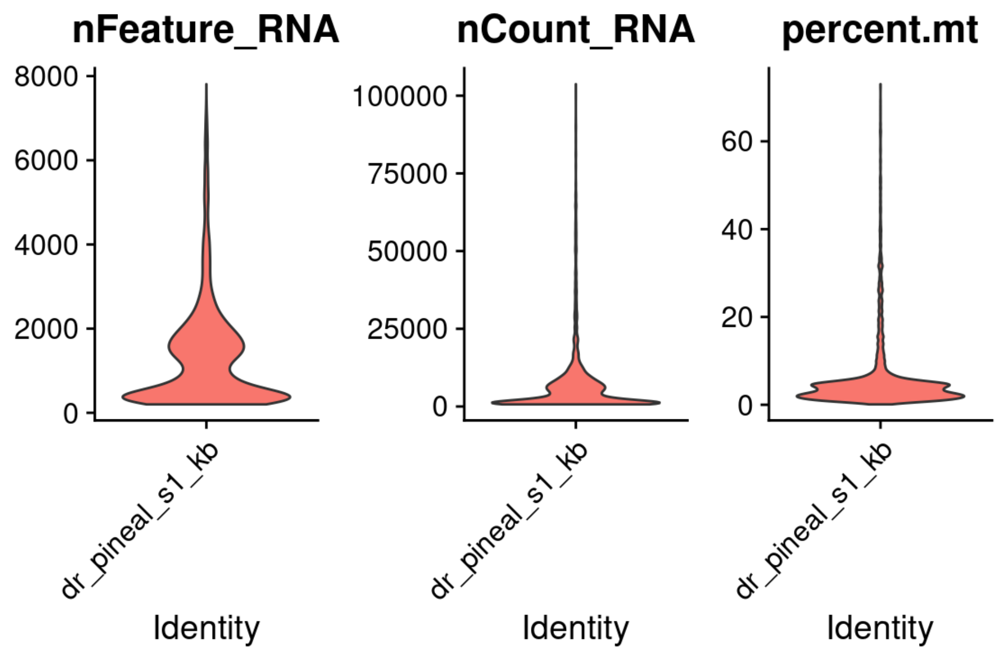


Total number of cells before filtration:

```
sum(table(...=pineal_s1_kb_forced_101@active.ident))
```

```
## [1] 4980
```

Filteration of outlier cells containing unusual number of genes, UMI or percentage of mitochondrial genes. Plot the distribution of the filtered cells.

```
pineal_s1_kb_forced_101 <- subset(x = pineal_s1_kb_forced_101,
                                  subset = nFeature_RNA > 200
                                  & nCount_RNA < 15000
                                  & percent.mt<30)

VlnPlot(object = pineal_s1_kb_forced_101,
        features = c("nFeature_RNA", "nCount_RNA", "percent.mt"),
        ncol = 3,
        pt.size=0)
```

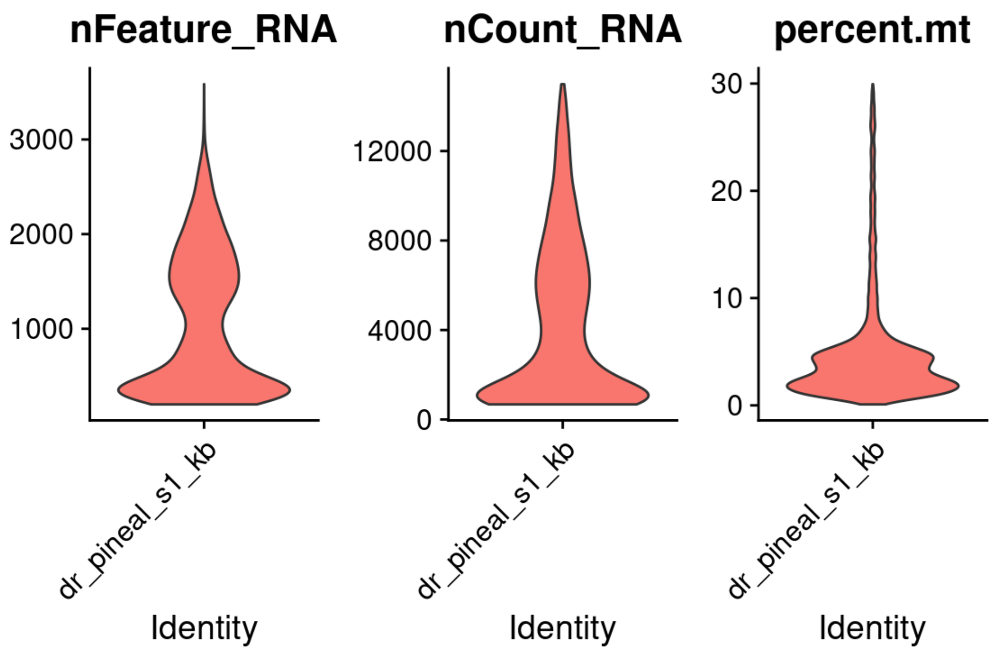


Total number of cells after filtration:

```
sum(table(...=pineal_s1_kb_forced_101@active.ident))
```

```
## [1] 4347
```

Standard normalization, variable gene identification and scaling:

```
pineal_s1_kb_forced_101 <- NormalizeData(object = pineal_s1_kb_forced_101,
                                         normalization.method = "LogNormalize",
                                         scale.factor = 10000)

pineal_s1_kb_forced_101 <- FindVariableFeatures(object = pineal_s1_kb_forced_101,
                                                selection.method = "vst",
                                                nfeatures = 2000)

all_genes_kallisto_forced_s1 <- rownames(x = pineal_s1_kb_forced_101)
pineal_s1_kb_forced_101 <- ScaleData(object = pineal_s1_kb_forced_101, features = all_genes_kall
isto_forced_s1)
```

Principal component analysis.

```
pineal_s1_kb_forced_101 <- RunPCA(object = pineal_s1_kb_forced_101, features = VariableFeatures
(object = pineal_s1_kb_forced_101))
```

```
## PC_ 1
## Positive:  selenop, krt8, zgc:162730, cd63, atp1b1a, junba, cxcl18b, s100a10b, si:ch73-335l2
1.4, atp1a1b
##     actb2, actb1, icn, zfp36l1b, krt18a.1, sparc, jdp2b, angptl4, fabp7a, fxyd1
##     tagln2, fosb, nfkbiaa, btg2, sgk1, CR318588.4, fosl1a, ctsla, ccn1, serpinh1b
## Negative:  rbp4l, pde6ha, gngt1, exorh, pde6ga, BX004774.2, pde6gb, gnat1, saga, tph2
##     rbp3, nrgna, cnga1a, rcvrn2, rcvrn3, rcvrnb, arl3l1, tulp1a, asmt, si:dkey-220f10.4
##     arl3l2, aipl2, ddc, cngb3.2, ppdpfa, cldn2, ahcy, CABZ01073265.1, parapinopsinb, si:ch211
-245j22.3
## PC_ 2
## Positive:  cx32.2, si:dkey-27i16.2, fcer1gl, cd74b, ctss2.2, cd74a, zgc:64051, ccl35.1, mhc2
a, lgals3bpb
##     si:cabz01074946.1, stoml3b, spi1b, havcr1, cxcr4b, slc7a7, ccl35.2, ccl34b.1, pfn1, ncf1
##     fermt3b, BX649485.1, ms4a17a.9, apoc1, arpc1b, cebpa, laptm5, lgals9l1, si:ch211-102c2.4,
ctsc
## Negative:  clu, atp1a1b, fabp7a, hspa8b, bckdhbl, stra6, hsp70l, scg3, cbx7a, si:ch211-81a5.8
##     vim, hspb8, krt18a.1, uchl1, cdkn1a, si:ch211-80h18.1, slc38a3a, slc13a1, wfdc2, s100b
##     dkk3b, proca1, si:dkey-11c5.11, mdkb, rgrb, zgc:114181, fabp7b, AL954359.2, rbp1.1, rdh5
## PC_ 3
## Positive:  rtn1b, elavl4, ywhag2, scg3, sncga, scg2b, syt1a, dusp5, si:dkey-33c12.3, snap25a
##     bdnf, ctsd, cbx7a, stmn4l, slc35g2a, vgf, chga, map1b, atp1a3a, stmn2a
##     id4, syngr3b, atp6v1b2, anxa13l, elavl3, si:ch73-119p20.1, uchl1, pcp4a, gng13b, gng3
## Negative:  hbaa1, hbba1.1, hbba1, si:ch211-5k11.8, hbba2, hbaa2, si:ch211-103n10.5, rbp4, ccl
25b, CABZ01092746.1
##     dcn, pmp22a, sost, igfbp5b, cldn11a, anxa1a, fsta, pcolcea, igfbp2a, lxn
##     bhmt, twist1a, thbs4b, msx1b, si:ch1073-291c23.2, clec3ba, eif4ebp3, serpinf1, vmo1a, si:
dkey-11f4.20
## PC_ 4
## Positive:  crhbp, si:dkey-33c12.3, chga, bdnf, elavl4, snap25a, vgf, elavl3, cxcl12a, anxa13l
##     id4, si:dkeyp-69c1.7, uts2a, stmn2a, stmn4l, nxph1, nell2b, gng13b, p2rx2, hpcal4
##     atp6v1b2, cart3, rtn1b, insm1b, nrsn1, atp1a3a, slc17a6b, syt1a, scg2b, pcp4a
## Negative:  si:ch211-81a5.8, ndrg1a, ppdpfa, bckdhbl, si:dkey-11c5.11, slc13a1, zgc:114181, cx
32.2, si:ch211-80h18.1, ahcy
##     stoml3b, dkk3b, pde6gb, fabp7b, cetn4, vim, rbp1.1, unc119.2, lgals3bpb, AL954359.2
##     arl3l2, gstm.3, ctss2.2, rbp3, nrgna, havcr1, spi1b, tulp1a, tmem98, UTP14C
## PC_ 5
## Positive:  slc13a1, dkk3b, zgc:114181, fabp7b, gstm.3, rbp1.1, bckdhbl, si:dkey-12l12.1, si:c
h211-80h18.1, mstnb
##     zgc:153311, tmem98, b3glcta, AL954359.2, si:dkey-11c5.11, flr, tmem72, slc38a3a, ahcyl1,
zgc:158404
##     bmp1b, tbata, vim, rbp2b, foxj1a, cyp19a1b, ackr3a, enkur, fgfbp1b.1, mctp2b
## Negative:  rdh5, asip2b, rgra, fxyd6l, rbp5, si:ch211-251b21.1, cxcl14, efhd1, rlbp1b, syt5b
##     c1qtnf5, jhy, slc6a2, tagln2, slc3a2a, ppp1r14aa, slc1a3a, coch, her15.2, cldn7a
##     scg3, fdx1, prdx1, cdon, myl9b, bzw1b, cnn2, slc1a2b, txn, marcksl1b
```

Visualize the principal components percentage of variance by an elbow plot.

```
ElbowPlot(object = pineal_s1_kb_forced_101, ndims = 30)
```

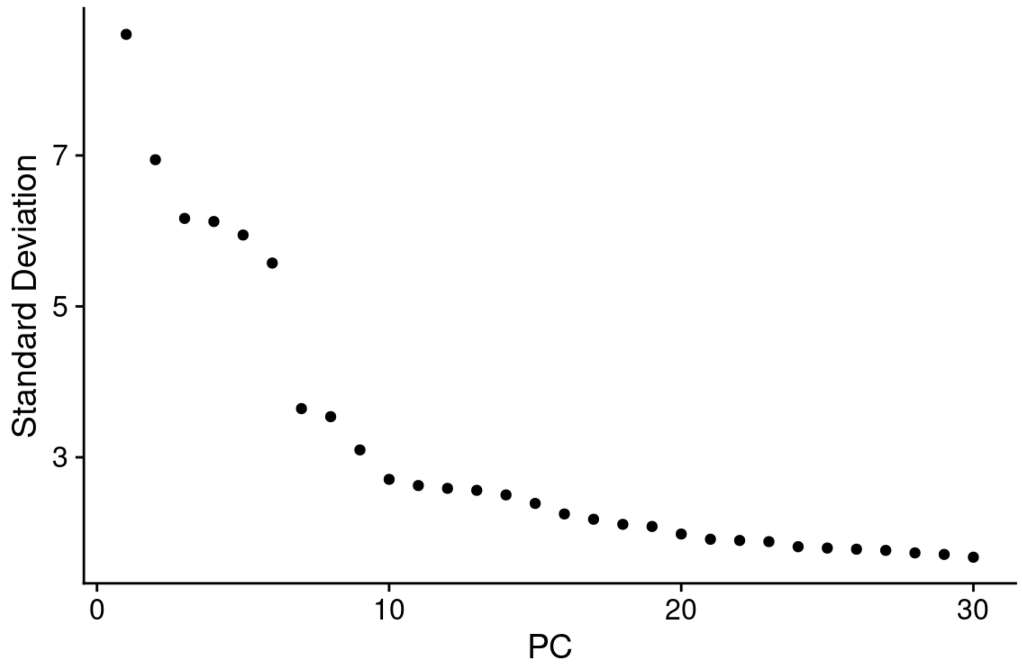


PCs 1-25 were used as dimensions of reduction to compute the k.param nearest neighbors

```
pineal_s1_kb_forced_101 <- FindNeighbors(object = pineal_s1_kb_forced_101, dims = 1:25)
pineal_s1_kb_forced_101 <- FindClusters(object = pineal_s1_kb_forced_101, resolution = 1.5)
```

```
## Modularity Optimizer version 1.3.0 by Ludo Waltman and Nees Jan van Eck
##
## Number of nodes: 4347
## Number of edges: 168165
##
## Running Louvain algorithm...
## Maximum modularity in 10 random starts: 0.8483
## Number of communities: 23
## Elapsed time: 0 seconds
```

```
pineal_s1_kb_forced_101 <- RunUMAP(object = pineal_s1_kb_forced_101, dims = 1:25)

kb_forced_UMAP_unmerged_s1_res_1_5 <- DimPlot(object = pineal_s1_kb_forced_101, reduction = "uma
p",
                            label=TRUE, pt.size = 0.5, label.size = 3) +
  theme(legend.position="none",
        axis.title.x=element_text(size=12),
        axis.title.y=element_text(size=12),
        plot.title = element_text(size=12, hjust=0.0)) + ggtitle("kallisto forced (res.=1.5)")
kb_forced_UMAP_unmerged_s1_res_1_5
```

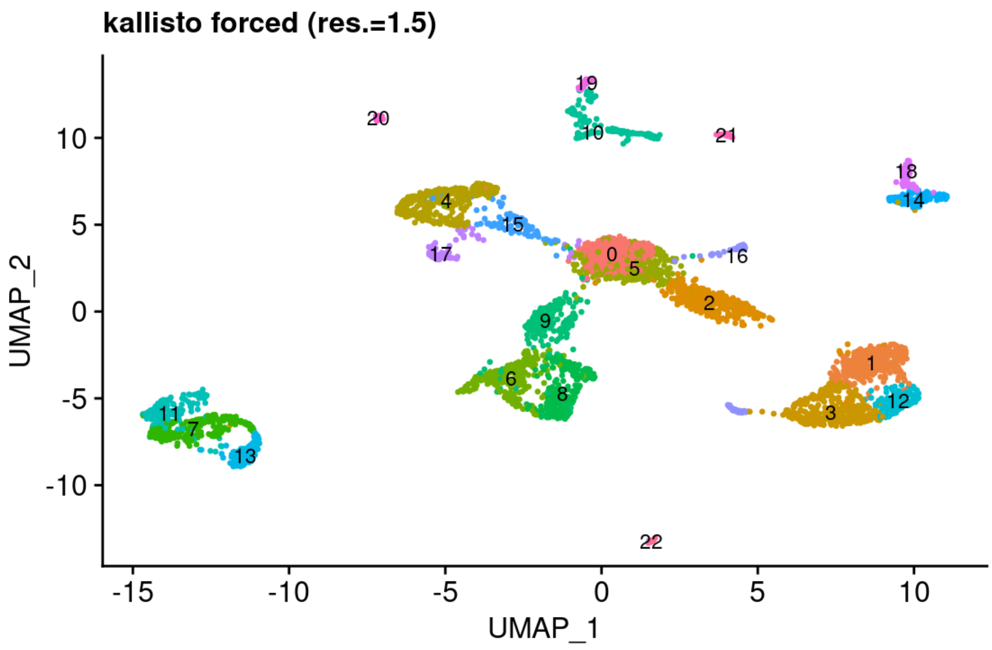


Analysis of the top markers for each cluster.

```
pineal_s1_kb_forced_101.markers <- FindAllMarkers(object = pineal_s1_kb_forced_101,
                                                  only.pos = TRUE,
                                                  min.pct = 0.25,
                                                  logfc.threshold = 0.8)
```

```
pineal_s1_kb_forced_101.markers %>% group_by(cluster) %>% top_n(n = 10, wt = avg_log2FC)
```

| p_val <dbl> | avg_log2FC <dbl> | pct.1 <dbl> | pct.2 <dbl> | p_val_adj <dbl> | cluster <fct> | gene <chr> |
|---|---|---|---|---|---|---|
| 0.000000e+00 | 3.376069 | 0.891 | 0.165 | 0.000000e+00 | 0 | cahz |
| 1.715462e-296 | 2.763641 | 0.927 | 0.216 | 3.854643e-292 | 0 | si:ch211-250g4.3 |
| 4.242644e-288 | 4.160237 | 1.000 | 0.995 | 9.533222e-284 | 0 | hbba1.1 |
| 3.824799e-286 | 4.435074 | 1.000 | 0.996 | 8.594324e-282 | 0 | hbaa1 |
| 5.095115e-286 | 4.146379 | 1.000 | 0.942 | 1.144872e-281 | 0 | hbba1 |
| 1.325773e-276 | 4.146113 | 1.000 | 0.908 | 2.979012e-272 | 0 | si:ch211-5k11.8 |
| 2.736486e-262 | 4.593392 | 0.994 | 0.826 | 6.148883e-258 | 0 | hbba2 |
| 8.820660e-253 | 2.979947 | 0.891 | 0.286 | 1.982002e-248 | 0 | si:ch211-103n10.5 |
| 1.912101e-245 | 4.228425 | 0.972 | 0.689 | 4.296490e-241 | 0 | hbaa2 |
| 6.337107e-195 | 2.333401 | 0.663 | 0.139 | 1.423948e-190 | 0 | nt5c2l1 |

1-10 of 230 rows Previous **1** 2 3 4 5 6 ... 23 Next

Dotplot of the top known markers of the pineal cell types (based on Shainer et al. 2019) as well as newly identify markers (such as dcn and ccr9a).

```
kallisto_forced_dotplot_unmerged_s1<- DotPlot(pineal_s1_kb_forced_101, features = dot_plot_genes
_s1,
                                        cluster.idents=FALSE, dot.scale=2) + RotatedAxis() +
  theme(axis.text.x = element_text(angle=45, size=10),
        axis.text.y = element_text(size=5, angle=0),
        legend.title = element_text(size=10),
        legend.text = element_text(size = 10),
        axis.title.x=element_blank(),
        axis.title.y=element_blank())
kallisto_forced_dotplot_unmerged_s1
```

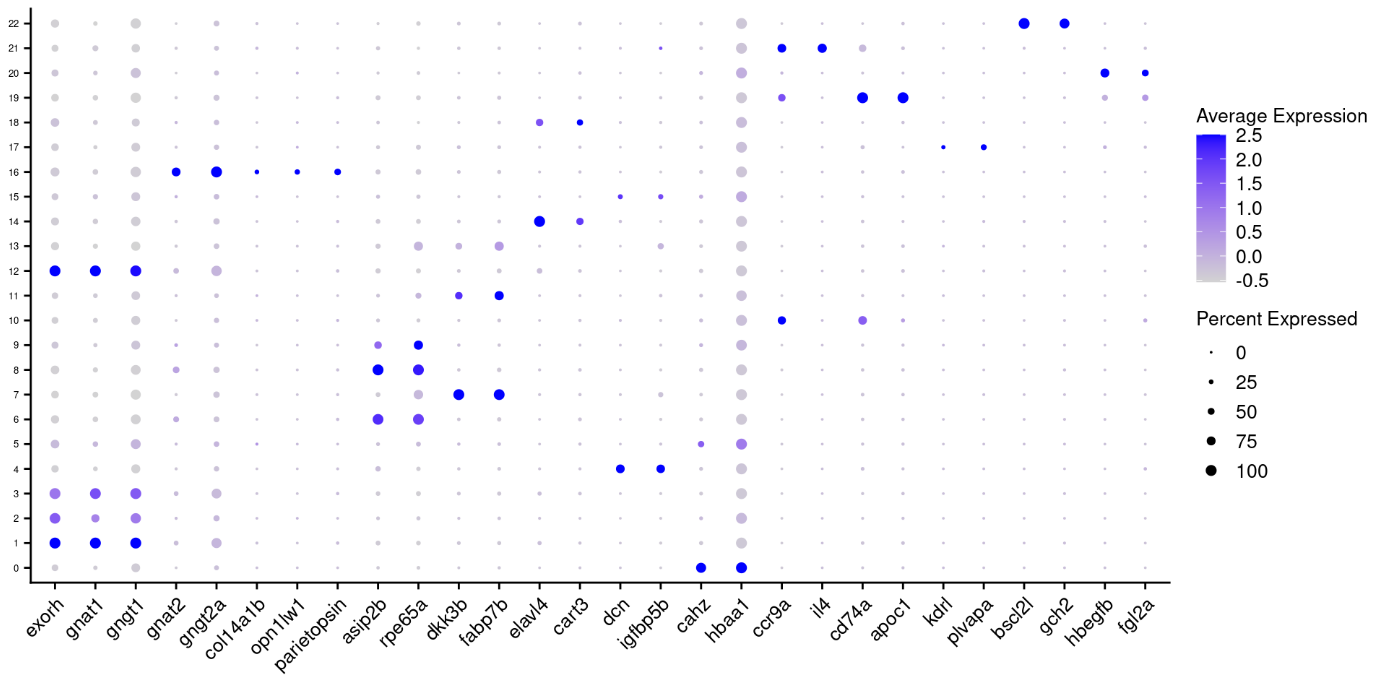


```
FeaturePlot(object=pineal_s1_kb_forced_101, features = c("opn1lw1", "parietopsin"), label = TRUE
, label.size = 3)
```

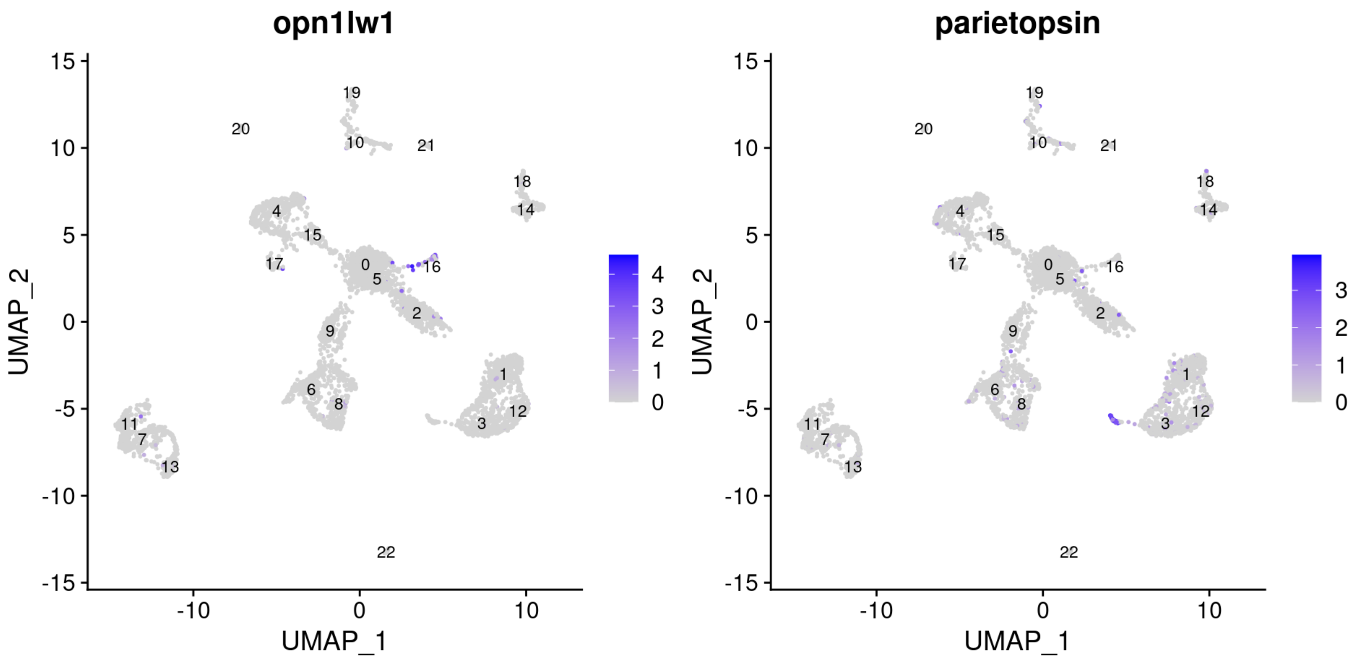


Under a resolution of 1.5, the green- and red-like photoreceptors cannot be separated. Increasing the resolution in the case of kallisto forced enable to seprate those photoreceptors.

```
pineal_s1_kb_forced_101 <- FindNeighbors(object = pineal_s1_kb_forced_101, dims = 1:25)
pineal_s1_kb_forced_101 <- FindClusters(object = pineal_s1_kb_forced_101, resolution = 2.4)
```

```
## Modularity Optimizer version 1.3.0 by Ludo Waltman and Nees Jan van Eck
##
## Number of nodes: 4347
## Number of edges: 168165
##
## Running Louvain algorithm...
## Maximum modularity in 10 random starts: 0.7926
## Number of communities: 30
## Elapsed time: 0 seconds
```

```
pineal_s1_kb_forced_101 <- RunUMAP(object = pineal_s1_kb_forced_101, dims = 1:25)

kb_forced_UMAP_unmerged_s1_res_2_4 <- DimPlot(object = pineal_s1_kb_forced_101, reduction = "uma
p",
                             label=TRUE, pt.size = 0.5, label.size = 3) +
  theme(legend.position="none",
        axis.title.x=element_text(size=12),
        axis.title.y=element_text(size=12),
        plot.title = element_text(size=14, hjust=0.0)) + ggtitle("kallisto forced (res.=2.4)") +
  theme(plot.title = element_text(size = 12))
kb_forced_UMAP_unmerged_s1_res_2_4
```

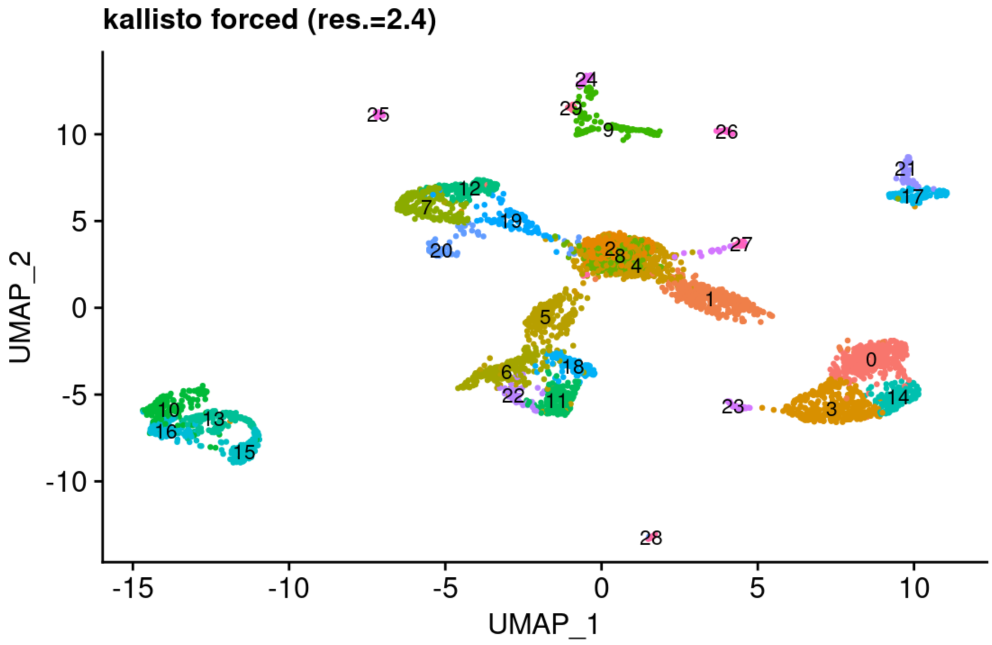


Dotplot high res

```
kallisto_forced_dotplot_hig_res_s1<- DotPlot(pineal_s1_kb_forced_101, features = dot_plot_genes_
s1,
                                         cluster.idents=FALSE, dot.scale=2) + RotatedAxis() +
  theme(axis.text.x = element_text(angle=45, size=10),
        axis.text.y = element_text(size=5, angle=0),
        legend.title = element_text(size=10),
        legend.text = element_text(size = 10),
        axis.title.x=element_blank(),
        axis.title.y=element_blank())
kallisto_forced_dotplot_hig_res_s1
```

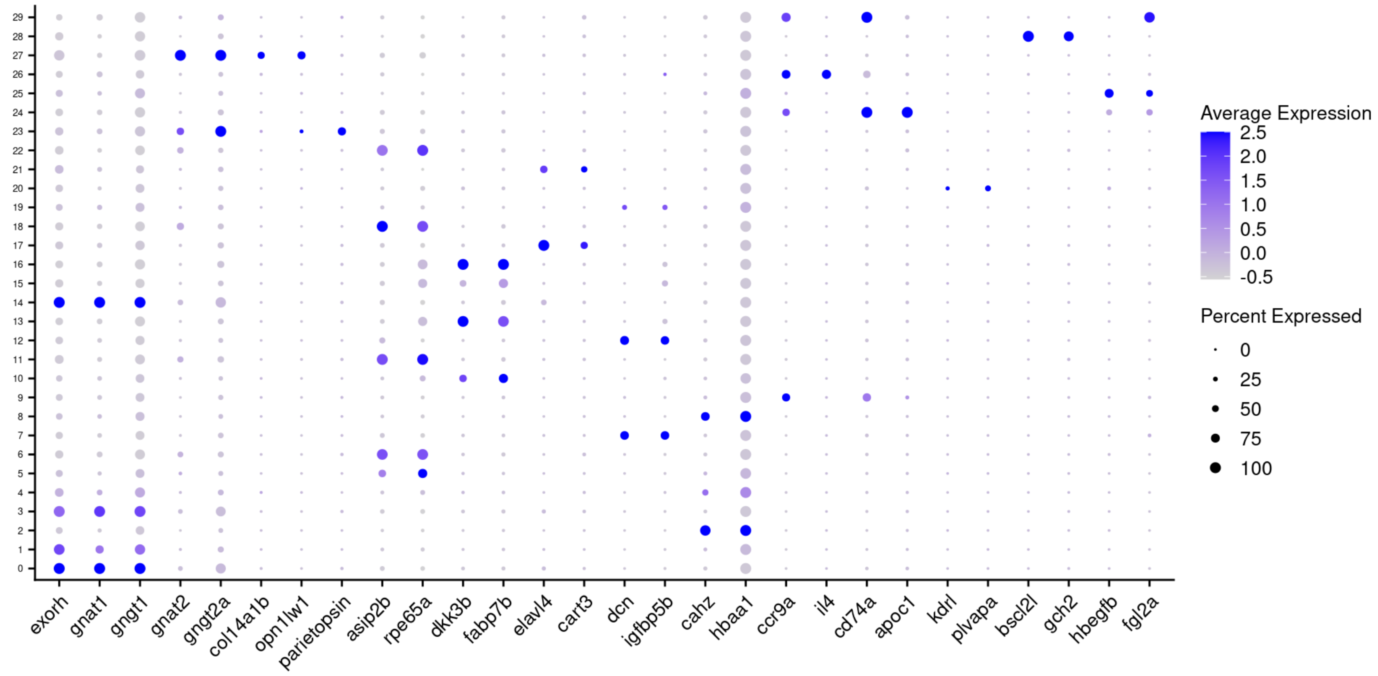


### plots

```
umap_S1<- ggarrange(cellr_UMAP_unmerged_s1,
                    cellr_UMAP_unmerged_s1_res_3_5,
                    kb_UMAP_unmerged_s1,
                    kb_forced_UMAP_unmerged_s1_res_1_5,
                    kb_forced_UMAP_unmerged_s1_res_2_4,
                  labels = c("A", "B", "C", "D","E"),
                  common.legend = FALSE,
                  ncol = 1, nrow = 5) #legend = "right")

dotplots_s1<- ggarrange(cellranger_dotplot_unmerged_s1,
                        cellranger_dotplot_high_res_s1,
                        kallisto_dotplot_unmerged_s1,
                        kallisto_forced_dotplot_unmerged_s1,
                        kallisto_forced_dotplot_hig_res_s1,
          common.legend = TRUE,
          ncol = 1, nrow = 5, legend = "right")

ggarrange(umap_S1, dotplots_s1,
          ncol = 2, nrow = 1, widths = c(1, 2))
```

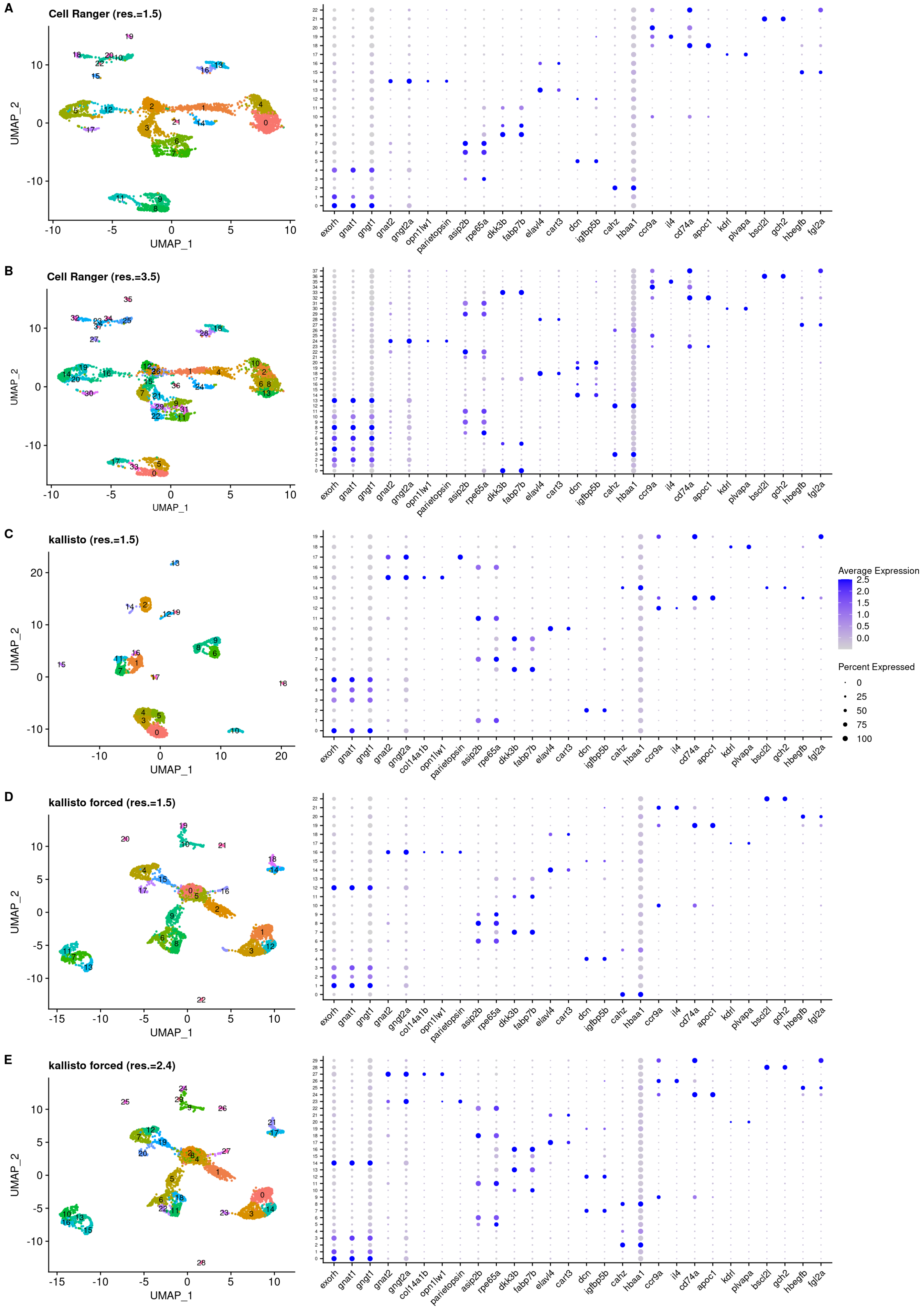
A
Cell Ranger (res.=1.5)
B
Cell Ranger (res.=3.5)
C
kallisto (res.=1.5)
D
kallisto forced (res.=1.5)
E
kallisto forced (res.=2.4)
UMAP_1
UMAP_2
exorh
gnat1
gngt1
gnat2
gngt2a
col14a1b
opn1lw1
parietopsin
asip2b
rpe65a
dkk3b
fabp7b
elavl4
cart3
dcn
igfbp5b
cahz
hbaa1
ccr9a
il4
cd74a
apoc1
kdrl
plvapa
bscl2l
gch2
hbegfb
fgl2a
Average Expression
2.5
2.0
1.5
1.0
0.5
0.0
Percent Expressed
0
25
50
75
100
