## Additional File 6 for "Choice of pre-processing pipeline influences clustering quality of scRNA-seq datasets"

---
title: "Zebrafish pineal gland scSeq sample 2 downstream analysis"
output:
  html_document:
    df_print: paged
  pdf_document: default
---

Analysis of zebrafish pineal gland cell types. For original data see Shainer et al. 2019 (<https://www.sciencedirect.com/science/article/pii/S0960982219305561>, pineal sample 2).
The RDS files were generated with the KB helper tool as described in the manuscript. This notebook demonstrate the downstream processing and cell type classification. The analysis is based on Seurat's standard workflow (<https://satijalab.org/seurat/articles/pbmc3k_tutorial.html>).

```{r message=FALSE, warning=FALSE}
library(dplyr)
library(Seurat)
library(Matrix)
library(ggplot2)
library(reticulate)
```

Load RDS files
```{r}
pineal_s2_cellr_101 <- readRDS("/zstorage/cellr_vs_kallisto/rds_files/D_rerio.GRCz11.101/dr_pineal_s2_cellr.rds")
pineal_s2_kb_101 <- readRDS("/zstorage/cellr_vs_kallisto/rds_files/D_rerio.GRCz11.101/dr_pineal_s2_kb.rds")
pineal_s2_kb_forced_101 <- readRDS("/zstorage/cellr_vs_kallisto/rds_files/D_rerio.GRCz11.101/dr_pineal_s2_kb_forced.rds")
```

## Downstream analysis of data preprocessed with kallisto

Calculate the percentage of mitochondrial genes per cell.
```{r}
pineal_s2_kb_101[["percent.mt"]] <- PercentageFeatureSet(object = pineal_s2_kb_101, pattern = "^mt-")
```

Visualize QC metrics.
```{r fig.height=4, fig.width=6}
VlnPlot(object = pineal_s2_kb_101,
        features = c("nFeature_RNA", "nCount_RNA", "percent.mt"),
        ncol = 3,
        pt.size=0)
```


Total number of cells before filtration:
```{r}
sum(table(...=pineal_s2_kb_101@active.ident))
```


Filtration of outlier cells containing unusual number of genes, UMI or percentage of mitochondrial genes. Plot the distribution of the filtered cells.
```{r fig.height=4, fig.width=6}
pineal_s2_kb_101 <- subset(x = pineal_s2_kb_101,
                           subset = nFeature_RNA > 200
                           & nCount_RNA < 15000
                           & percent.mt<30)
VlnPlot(object = pineal_s2_kb_101,
        features = c("nFeature_RNA", "nCount_RNA", "percent.mt"),
        ncol = 3,
        pt.size=0)
```


Total number of cells after filtration:
```{r}
sum(table(...=pineal_s2_kb_101@active.ident))
```


Standard normalization, variable gene identification and scaling:
```{r echo=TRUE, message=FALSE, warning=FALSE}
pineal_s2_kb_101 <- NormalizeData(object = pineal_s2_kb_101,
                                  normalization.method = "LogNormalize",
                                  scale.factor = 10000)

pineal_s2_kb_101 <- FindVariableFeatures(object = pineal_s2_kb_101,
                                         selection.method = "vst",
                                         nfeatures = 2000)

all_genes_kallisto <- rownames(x = pineal_s2_kb_101)
pineal_s2_kb_101 <- ScaleData(object = pineal_s2_kb_101,
                              features = all_genes_kallisto)

```


Principal component analysis.
```{r}
pineal_s2_kb_101 <- RunPCA(object = pineal_s2_kb_101,
                           features = VariableFeatures(object = pineal_s2_kb_101))
```


Visualize the principal components percentage of variance by an elbow plot.
```{r fig.height=4, fig.width=6}
ElbowPlot(object = pineal_s2_kb_101, ndims = 30)
```


PCs 1-20 were used as dimensions of reduction to compute the k.param nearest neighbors
```{r echo=TRUE, message=FALSE, warning=FALSE}
pineal_s2_kb_101 <- FindNeighbors(object = pineal_s2_kb_101, dims = 1:20)
pineal_s2_kb_101 <- FindClusters(object = pineal_s2_kb_101, resolution = 0.9)

#Run non-linear dimensional reduction (UMAP)
pineal_s2_kb_101 <- RunUMAP(object = pineal_s2_kb_101, dims = 1:20)
```
```{r fig.height=4, fig.width=6}
kb_UMAP_unmerged <- DimPlot(object = pineal_s2_kb_101, reduction = "umap",
                            label=TRUE, pt.size = 0.5, label.size = 3) +
  theme(legend.position="none",
        axis.title.x=element_text(size=10),
        axis.title.y=element_text(size=10),
        plot.title = element_text(size=14, hjust=0.0)) + ggtitle("kallisto")
kb_UMAP_unmerged
```


Analysis of the top markers for each cluster.
```{r echo=TRUE, message=FALSE, warning=FALSE}
pineal_s2_kb_101.markers <- FindAllMarkers(object = pineal_s2_kb_101,
                                           only.pos = TRUE,
                                           min.pct = 0.25,
                                           logfc.threshold = 0.8)
```
```{r}
pineal_s2_kb_101.markers %>% group_by(cluster) %>% top_n(n = 10, wt = avg_log2FC)
```

Dotplot of the top known markers of the pineal cell types (based on Shainer et al. 2019) as well as newly identify markers (such as col14a1b, dcn, ccr9a and epcan).

```{r fig.height=4, fig.width=6}
dot_plot_genes= c("exorh", "gnat1", "gngt1",
                  "gnat2", "gngt2a", "col14a1b", "opn1lw1","parietopsin",
                  "asip2b", "rpe65a",
                  "dkk3b", "fabp7b",
                  "elavl4", "cart3",
                  "gng8", "kiss1",
                  "dcn", "igfbp5b",
                  "cahz", "hbaa1",
                  "ccr9a", "il4",
                  "cd74a", "apoc1",
                  "kdrl", "plvapa",
                  "epcam", "icn2")

```
```{r fig.height=5, fig.width=10}
kallisto_dotplot_unmerged<- DotPlot(pineal_s2_kb_101, features = dot_plot_genes, cluster.idents=FALSE, dot.scale=4) + RotatedAxis() +
  theme(axis.text.x = element_text(angle=45, size=10),
        axis.text.y = element_text(size=5, angle=0),
        legend.title = element_text(size=10),
        legend.text = element_text(size = 10),
        axis.title.x=element_blank(),
        axis.title.y=element_blank()) + ggtitle("kallisto")
kallisto_dotplot_unmerged
```


Cell type identity was assigned based on the previous analysis of the pineal cell types (Shainer et al. 2019), while newly detected clusters were identified by comparing their marker genes to what is known in literature. The clustering analysis resulted in sub-clusters of the known pineal clusters (clusters 1,2,3,7 &21 are the rod-like PhRs, 0, 9 & 17 are the RPE-like and 11 & 12 are the Muller glia-like) as well as the habenula kiss1 neurons (clusters 4,5, & 20) and the leukocytes (clusters 13 & 22). The sub-clusters which had similar markers, varying only in expression levels, were merged to simplify the visualization.

```{r}
new.kb.cluster.ids <- c("RPE-like",
                     "rod-like PhR",
                     "rod-like PhR",
                     "rod-like PhR",
                     "habanula neurons (kiss1)",
                     "habanula neurons (kiss1)",
                     "fibroblasts",
                     "rod-like PhR",
                     "habanula neurons (gng8)",
                     "RPE-like",
                     "erythrocytes",
                     "muller glia-like",
                     "muller glia-like",
                     "leukocytes",
                     "neurons",
                     "microglia/macrophages",
                     "red cone-like PhR",
                     "RPE-like",
                     "green cone-like PhR",
                     "vascular endothelial cells",
                     "habanula neurons (kiss1)",
                     "rod-like PhR",
                     "leukocytes",
                     "epithelial cells")
names(new.kb.cluster.ids) <- levels(pineal_s2_kb_101)
pineal_s2_kb_101 <- RenameIdents(pineal_s2_kb_101, new.kb.cluster.ids)
```
```{r fig.height=4, fig.width=8}
kallisto_umap<- DimPlot(object = pineal_s2_kb_101, reduction = "umap",
        label=TRUE, pt.size = 0.5)+ theme(axis.title.x=element_text(size=12),
                                          axis.title.y=element_text(size=12),
                                          plot.title = element_text(size=14, hjust=0.0)) + ggtitle("kallisto")
kallisto_umap
```
```{r}
```


New dotplot (arrange clusters in a common order)
```{r}
library(forcats)
pineal_s2_kb_101@active.ident <- fct_relevel(pineal_s2_kb_101@active.ident,
                                             "rod-like PhR",
                                             "red cone-like PhR",
                                             "green cone-like PhR",
                                             "RPE-like","muller glia-like",
                                             "neurons", "habanula neurons (gng8)",
                                             "habanula neurons (kiss1)","fibroblasts",
                                             "erythrocytes", "leukocytes",
                                             "microglia/macrophages",
                                             "vascular endothelial cells",
                                             "epithelial cells")
```
```{r fig.height=5, fig.width=10}
kallisto_dotplot<- DotPlot(pineal_s2_kb_101, features = dot_plot_genes, cluster.idents=FALSE, dot.scale=4) + RotatedAxis() +
  theme(axis.text.x = element_text(angle=45, size=10),
        axis.text.y = element_text(size=12, angle=0),
        legend.title = element_text(size=12),
        legend.text = element_text(size = 12),
        axis.title.x=element_blank(),
        axis.title.y=element_blank()) + ggtitle("kallisto")
kallisto_dotplot
```

How many cells in each cluster?
```{r}
(table(...=pineal_s2_kb_101@active.ident))
```


## Downstream analysis of data preprocessed with Cell Ranger

Calculate percentage of mitochondrial genes per cell and visualize QC metrics
```{r fig.height=4, fig.width=6}
pineal_s2_cellr_101[["percent.mt"]] <- PercentageFeatureSet(object = pineal_s2_cellr_101, pattern = "^mt-")
VlnPlot(object = pineal_s2_cellr_101, features = c("nFeature_RNA", "nCount_RNA", "percent.mt"), ncol = 3., pt.size=0)
```


Total number of cells before filtration:
```{r}
sum(table(...=pineal_s2_cellr_101@active.ident))
```


Filteration of outlier cells containing unusual number of genes, UMI or percentage of mitochondrial genes. Plot the distributation of the filtered cells and print the total number of cells.
```{r fig.height=4, fig.width=6}
pineal_s2_cellr_101 <- subset(x = pineal_s2_cellr_101, subset = nFeature_RNA > 200 &  nCount_RNA < 15000 & percent.mt<30)
VlnPlot(object = pineal_s2_cellr_101, features = c("nFeature_RNA", "nCount_RNA", "percent.mt"), ncol = 3., pt.size=0)

```


Total number of cells after filtration:
```{r}
sum(table(...=pineal_s2_cellr_101@active.ident))
```


Standard normalization, variable gene identification and scaling:
```{r echo=TRUE, message=FALSE, warning=FALSE}
pineal_s2_cellr_101 <- NormalizeData(object = pineal_s2_cellr_101, normalization.method = "LogNormalize", scale.factor = 10000)

pineal_s2_cellr_101 <- FindVariableFeatures(object = pineal_s2_cellr_101, selection.method = "vst", nfeatures = 2000)

all_genes_cellr <- rownames(x = pineal_s2_cellr_101)
pineal_s2_cellr_101 <- ScaleData(object = pineal_s2_cellr_101, features = all_genes_cellr)
```


Principal component analysis.
```{r}
pineal_s2_cellr_101 <- RunPCA(object = pineal_s2_cellr_101, features = VariableFeatures(object = pineal_s2_cellr_101))
```


Visualize the principal components percentage of variance by an elbow plot.
```{r fig.height=4, fig.width=6}
ElbowPlot(object = pineal_s2_cellr_101, ndims = 30)
```


PCs 1-20 were used as dimensions of reduction to compute the k.param nearest neighbors
```{r echo=TRUE, message=FALSE, warning=FALSE}
pineal_s2_cellr_101 <- FindNeighbors(object = pineal_s2_cellr_101, dims = 1:20)
pineal_s2_cellr_101 <- FindClusters(object = pineal_s2_cellr_101, resolution = 0.9)
#Run non-linear dimensional reduction (UMAP)
pineal_s2_cellr_101 <- RunUMAP(object = pineal_s2_cellr_101, dims = 1:20)
```
```{r fig.height=4, fig.width=6}
cr_UMAP_unmerged <- DimPlot(object = pineal_s2_cellr_101, reduction = "umap",
                            label=TRUE, pt.size = 0.5, label.size = 3) +
  theme(legend.position="none",
      axis.title.x=element_text(size=10),
      axis.title.y=element_text(size=10),
      plot.title = element_text(size=14, hjust=0.0)) + ggtitle("Cell Ranger")
cr_UMAP_unmerged
```


Analysis of the top markers for each cluster
```{r echo=TRUE, message=FALSE, warning=FALSE}
pineal_s2_cellr_101.markers <- FindAllMarkers(object = pineal_s2_cellr_101,
                                              only.pos = TRUE,
                                              min.pct = 0.25,
                                              logfc.threshold = 0.8)
```
```{r}
pineal_s2_cellr_101.markers %>% group_by(cluster) %>% top_n(n = 10, wt = avg_log2FC)
```


Plotting marker genes
```{r fig.height=5, fig.width=10}
cellr_dotplot_unmerged<- DotPlot(pineal_s2_cellr_101, features = dot_plot_genes, cluster.idents=FALSE, dot.scale=4) + RotatedAxis() +
  theme(axis.text.x = element_text(angle=45, size=10),
        axis.text.y = element_text(size=5, angle=0),
        legend.title = element_text(size=10),
        legend.text = element_text(size = 10),
        axis.title.x=element_blank(),
        axis.title.y=element_blank()) + ggtitle("Cell Ranger")
cellr_dotplot_unmerged
```


As before, the resulting over-clustering of the known pineal clusters (clusters 0, 1, 8 & 18 which are the rod-like PhRs, clusters 2, 5 & 16 which are the RPE-like and clusters 7 & 12  which are the Muller glia-like) as well as the habenula kiss1 neurons (clusters 3 & 10) and the leukocytes (clusters 11 & 19) were merged to simplify the visualization.

```{r}
#Rename the clusters
new.cr.cluster.ids <- c("rod-like PhR",
                        "rod-like PhR",
                        "RPE-like",
                        "habanula neurons (kiss1)",
                        "erythrocytes",
                        "RPE-like",
                        "fibroblasts",
                        "muller glia-like",
                        "rod-like PhR",
                        "habanula neurons (gng8)",
                        "habanula neurons (kiss1)",
                        "leukocytes",
                        "muller glia-like",
                        "neurons",
                        "cone-like PhR",
                        "microglia/macrophages",
                        "RPE-like",
                        "vascular endothelial cells",
                        "rod-like PhR",
                        "leukocytes",
                        "epithelial cells")
names(new.cr.cluster.ids) <- levels(pineal_s2_cellr_101)
pineal_s2_cellr_101 <- RenameIdents(pineal_s2_cellr_101, new.cr.cluster.ids)
```
```{r fig.height=4, fig.width=8}
cellranger_umap<- DimPlot(object = pineal_s2_cellr_101, reduction = "umap",
        label=TRUE, pt.size = 0.5) + theme(axis.title.x=element_text(size=12),
                                            axis.title.y=element_text(size=12),
                                            plot.title = element_text(size=14, hjust=0.0)) + ggtitle("Cell Ranger")

cellranger_umap
```


New dotplot (arrange clusters in a common order)
```{r}
pineal_s2_cellr_101@active.ident <- fct_relevel(pineal_s2_cellr_101@active.ident,
                                                "rod-like PhR", "cone-like PhR", "RPE-like",
                                                "muller glia-like", "neurons",
                                                "habanula neurons (gng8)", "habanula neurons (kiss1)",
                                                "fibroblasts", "erythrocytes", "leukocytes",
                                                "microglia/macrophages", "vascular endothelial cells",
                                                "epithelial cells")

```
```{r fig.height=5, fig.width=10}
cellranger_dotplot<- DotPlot(pineal_s2_cellr_101, features = dot_plot_genes, cluster.idents=FALSE, dot.scale=4) + RotatedAxis() +
  theme(axis.text.x = element_text(angle=45, size=10),
        axis.text.y = element_text(size=12, angle=0),
        legend.title = element_text(size=12),
        legend.text = element_text(size = 12),
        axis.title.x=element_blank(),
        axis.title.y=element_blank()) + ggtitle("Cell Ranger")
cellranger_dotplot
```

How many cells in each cluster?
```{r}
table(...=pineal_s2_cellr_101@active.ident)
```


## Downstream analysis of data preprocessed with kallisto forced

Similar process as before.
```{r fig.height=4, fig.width=6}
pineal_s2_kb_forced_101[["percent.mt"]] <- PercentageFeatureSet(object = pineal_s2_kb_forced_101, pattern = "^mt-")
VlnPlot(object = pineal_s2_kb_forced_101,
        features = c("nFeature_RNA", "nCount_RNA", "percent.mt"),
        ncol = 3,
        pt.size=0)
```


Total number of cells before filtering
```{r}
table(...=pineal_s2_kb_forced_101@active.ident)
```
```{r fig.height=4, fig.width=6}
pineal_s2_kb_forced_101 <- subset(x = pineal_s2_kb_forced_101, subset = nFeature_RNA > 200 &  nCount_RNA < 15000 & percent.mt<30)
VlnPlot(object = pineal_s2_kb_forced_101,
        features = c("nFeature_RNA", "nCount_RNA", "percent.mt"),
        ncol = 3,
        pt.size=0)
```


Total number of cells after filtering
```{r}
table(...=pineal_s2_kb_forced_101@active.ident)
```

```{r fig.height=4, fig.width=6, message=FALSE, warning=FALSE}
pineal_s2_kb_forced_101 <- NormalizeData(object = pineal_s2_kb_forced_101,
                                         normalization.method = "LogNormalize",
                                         scale.factor = 10000)

pineal_s2_kb_forced_101 <- FindVariableFeatures(object = pineal_s2_kb_forced_101,
                                                selection.method = "vst",
                                                nfeatures = 2000)

all_genes_kallisto_forced <- rownames(x = pineal_s2_kb_forced_101)
pineal_s2_kb_forced_101 <- ScaleData(object = pineal_s2_kb_forced_101,
                                     features = all_genes_kallisto_forced)

pineal_s2_kb_forced_101 <- RunPCA(object = pineal_s2_kb_forced_101,
                                  features = VariableFeatures(object = pineal_s2_kb_forced_101))
ElbowPlot(object = pineal_s2_kb_forced_101, ndims = 30)
```

```{r fig.height=4, fig.width=6}
pineal_s2_kb_forced_101 <- FindNeighbors(object = pineal_s2_kb_forced_101, dims = 1:20)
pineal_s2_kb_forced_101 <- FindClusters(object = pineal_s2_kb_forced_101, resolution = 1.2)
pineal_s2_kb_forced_101 <- RunUMAP(object = pineal_s2_kb_forced_101, dims = 1:20)
kb_forced_UMAP_unmerged <- DimPlot(object = pineal_s2_kb_forced_101, reduction = "umap",
                            label=TRUE, pt.size = 0.5, label.size = 3) +
  theme(legend.position="none",
        axis.title.x=element_text(size=10),
        axis.title.y=element_text(size=10),
        plot.title = element_text(size=14, hjust=0.0)) + ggtitle("kallisto forced")
kb_forced_UMAP_unmerged
```

```{r fig.height=5, fig.width=10}
kb_forced_dotplot_unmerged<- DotPlot(pineal_s2_kb_forced_101, features = dot_plot_genes, cluster.idents=FALSE, dot.scale=4) + RotatedAxis() +
  theme(axis.text.x = element_text(angle=45, size=10),
        axis.text.y = element_text(size=5, angle=0),
        legend.title = element_text(size=10),
        legend.text = element_text(size = 10),
        axis.title.x=element_blank(),
        axis.title.y=element_blank()) + ggtitle("kallisto forced")
kb_forced_dotplot_unmerged
```

```{r echo=TRUE, message=FALSE, warning=FALSE}
pineal_s2_kb_forced_101.markers <- FindAllMarkers(object = pineal_s2_kb_forced_101,
                                                  only.pos = TRUE,
                                                  min.pct = 0.25,
                                                  logfc.threshold = 0.8)
```
```{r}
pineal_s2_kb_forced_101.markers %>% group_by(cluster) %>% top_n(n = 10, wt = avg_log2FC)
```


## Total counts comparison

This part of the code is designed to analyze the differences in gene counts between the pineal sample 2 data that was pre-processed with Cell Ranger vs. kallisto.

```{r}
#load the required packages
library(pheatmap)
library(RColorBrewer)
library(viridisLite)
```

First we generate a data frame containing all the genes and their total counts. This data frame is based on the gene-cell matrix of the scSeq (the values of each row are summed up).

```{r}
# create a data frame with the total counts of cellranger
#copy the gene-cell matrix values:
cellr_counts<-data.frame(pineal_s2_cellr_101@assays[["RNA"]]@counts)
#sum the values of each row:
cellr_counts_row_sum<-data.frame(rowSums(cellr_counts))
#copy the gene names to a new column
cellr_counts_row_sum$gene<-row.names(cellr_counts_row_sum)
# change to row names to numbers, length equalr to the length of the gene-cell matrix.
row.names(cellr_counts_row_sum) <- 1:20701

# create a data frame with the total counts of kallisto (same steps as before)
kb_counts<-data.frame(pineal_s2_kb_101@assays[["RNA"]]@counts)
kb_counts_row_sum<-data.frame(rowSums(kb_counts))
kb_counts_row_sum$gene<-row.names(kb_counts_row_sum)
row.names(kb_counts_row_sum) <- 1:23387
```


Join the two dataframes (kallisto and Cell Ranger total counts) into one called "all counts".
Then, calculate the diff_ratio: (kb_counts - cellr_count)/(kb_counts + cellr_counts), which represents the count difference ratio.
Positive values are genes with higher counts in kallisto, negative values are genes with higher counts in Cell Ranger.
Values equal to 1 represent genes that are not detected at all in Cell Ranger (counts = 0),
and even if are counted only once in kallisto, would have the highest diff_ratio.
Similarly, values equal to -1 represent genes that are not detected at all in kallisto (counts = 0),
and even if are counted only once in Cell Ranger, would have the lowest diff_ratio.
We will remove those cases (diff_ratio=1 or diff_ratio=-1) later, in order to represent only genes truly detected with a reasonable number of counts.
For some genes that are not detected at all in one data set, the count value is "NA". Those genes, even if are highly counted in one dataset, will not have a diff_ratio value, but a "NA" instead. We will add the genes with the highest counts in one dataset, and "NA" in the other, for the downstream analyses as well, as those represent true diff_ratio cases, even though the "NA" might prevent them from passing the threshold.

```{r}
#join the two dataframes
all_counts<-full_join(kb_counts_row_sum, cellr_counts_row_sum, by = "gene")
#reorder the columns
all_counts <- all_counts[, c(2, 1, 3)]
# add a new column containg the dif ratio (as described above).
all_counts<-mutate(all_counts, diff_ratio = ((all_counts$rowSums.kb_counts.- all_counts$rowSums.cellr_counts.)/(all_counts$rowSums.kb_counts. + all_counts$rowSums.cellr_counts.)))
#name the columns currectly
names(all_counts)[2]="kallisto_counts"
names(all_counts)[3]="cellranger_counts"
# save the results in a csv file
write.csv(all_counts, "all_counts.csv", row.names = FALSE)
```

### Genes with higher counts in kallisto
Now we pick the top 80 diff_ratio genes in kallisto (diff_ratio closest to 1, but not equal to 1)
```{r}
#create a table of kb highly expressed genes that are very low in cellr. We first take the
top_80_diff_kb<- all_counts[all_counts$diff_ratio != 1, ] %>% top_n(n = 80, wt = diff_ratio)
#keep only the gene names
top_80_diff_kb<-data.frame(top_80_diff_kb$gene)
names(top_80_diff_kb)[1]="gene"
```


add the top 20 genes that exist in kallisto and not at all in Cell Ranger (assigned as NA)
```{r}
# find genes that only exist in kallisto
NA_genes_in_cellr<-setdiff(kb_counts_row_sum$gene,cellr_counts_row_sum$gene)
# use this dataframe to filter the NA rows from the diff_ratio table
NA_genes_in_cellr_counts_in_kb<-data.frame(filter(kb_counts_row_sum, kb_counts_row_sum$gene %in% NA_genes_in_cellr))
#pick the top 20 genes with high counts in kallisto and "NA" in cellranger
top_20_NA_kb<-NA_genes_in_cellr_counts_in_kb %>% top_n(n = 20, wt = rowSums.kb_counts.)
top_20_NA_kb<-data.frame(top_20_NA_kb$gene)
names(top_20_NA_kb)[1]="gene"
```

Now we combine the top 80 diff_ratio and the 20 NA genes
```{r}
top100_heatmap_kb<-bind_rows(top_80_diff_kb, top_20_NA_kb)
```

We want to plot those genes in a heatmap to see whether they are considered to be cluster markers
(and therefore their detection in a certain pre-processed dataset is the reason for the detection of unique cell type).
First we average the expression of the 100 genes from before, for each of the clusters. We average the scale.data.
```{r}
kb_RNA_heatmap<- as.data.frame(AverageExpression(
  pineal_s2_kb_101,
  return.seurat = FALSE,
  group.by = "ident",
  slot = "scale.data",
  verbose = TRUE,
  features=top100_heatmap_kb$gene
))

```

Now we can plot this data in a heatmap
```{r}
pheatmap(kb_RNA_heatmap,
         scale = "none",
         main= "High count genes in kb and their cluster specific average expression",
         color= viridis(11),
         border_color=NA,
         cluster_rows = TRUE,
         cluster_cols = FALSE,
         show_colnames = TRUE,
         angle_col=45,
         fontsize = 8, fontsize_row = 4)
```

### Genes with higher counts in Cell Ranger

Create a table of Cell Ranger highly expressed genes that are very low in kallisto (as before).
First,we pick the top 80 diff_ratio genes in Cell Ranger (diff_ratio closest to -1, but not equal to -1)

```{r}
top_80_diff_cellr<- all_counts[all_counts$diff_ratio != -1, ] %>% top_n(n = 80, wt = diff_ratio)
#keep only the gene names
top_80_diff_cellr<-data.frame(top_80_diff_cellr$gene)
names(top_80_diff_cellr)[1]="gene"
```

Add the top 20 genes that exist in Cell Ranger and not at all in kallisto (assigned as NA)
```{r}
# find genes that only exist in cellranger
NA_genes_in_kb<-setdiff(cellr_counts_row_sum$gene, kb_counts_row_sum$gene)
# use this dataframe to filter the NA rows from the diff_ratio table
NA_genes_in_kb_counts_in_cellr<-data.frame(filter(cellr_counts_row_sum, cellr_counts_row_sum$gene %in% NA_genes_in_kb))
#pick the top 20 genes with high counts in cellranger and "NA" in kallisto
top_20_NA_cellr<-NA_genes_in_kb_counts_in_cellr %>% top_n(n = 20, wt = rowSums.cellr_counts.)
top_20_NA_cellr<-data.frame(top_20_NA_cellr$gene)
names(top_20_NA_cellr)[1]="gene"
```


Now we combine the top 80 diff_ratio and the 20 NA genes
```{r}
top100_heatmap_cellr<-bind_rows(top_80_diff_cellr, top_20_NA_cellr)
```

We want to plot those genes in a heatmap to see whether they are considered to be cluster markers
(and therefore their detection in a certain pre-processed dataset is the reason for the detection of unique cell type).
First we average the expression of the 100 genes from before, for each of the clusters. We average the scale.data.

```{r}
cellr_RNA_heatmap<- as.data.frame(AverageExpression(
  pineal_s2_cellr_101,
  return.seurat = FALSE,
  group.by = "ident",
  slot = "scale.data",
  verbose = TRUE,
  features=top100_heatmap_cellr$gene
))
```

Now we can plot this data in a heatmap.
```{r}
pheatmap(cellr_RNA_heatmap,
         scale = "none",
         main= "High count genes in cellranger and their cluster specific average expression",
         color= viridis(11),
         border_color=NA,
         cluster_rows = TRUE,
         cluster_cols = FALSE,
         show_colnames = TRUE,
         angle_col=45,
         fontsize = 8, fontsize_row = 4)
```
